# Temporal RNA splicing dysregulation in a human multicellular model of *C9orf72* ALS

**DOI:** 10.64898/2026.10.13.755334

**Authors:** Berna Aliya, Shiqi Liu, Alison Mansfield, Solan Xiyuan Yao, Sabra Mouhi, Supriya S. Wariyar, Jesse Seth Tenkorang, Nuri E. Smith, Mingee Chung, Ah Joung Rachel Yu, Taylor Pio, Najah E.J. Smith, Sabrina H. Zhu, Adam N. Trautwig, Janani Parameswaran, Luming Yin, Caroline M. Watson, Brain Organoid Hub, Jie Jiang, Nicholas T. Seyfried, Zachary T. McEachin, Bing Yao, Jimena Andersen

## Abstract

Amyotrophic lateral sclerosis (ALS) is a complex and heterogenous disease, comprising of familial and sporadic forms, numerous genetic mutations, and a wide range of affected cell types and proposed dysregulated processes. While the usage of *in vitro* cell cultures, mouse models, and postmortem patient tissue have advanced our understanding of ALS disease mechanisms, the temporal progression of the disease and how early molecular changes evolve remain poorly understood. Here, we establish human spinal cord organoids (hSpO) derived from *C9orf72* patient induced pluripotent stem cells (C9-ALS hiPSCs) as a platform to model longitudinal changes across the disease trajectory. We show that C9-ALS hSpO recapitulate hallmark *C9orf72*-related pathologies, increased cell death, and progressive neuronal dysfunction. We further identify RNA splicing dysregulation as an early and persistent feature of C9-ALS, with splicing signatures in hSpO closely mirroring those observed in postmortem patient tissue. Finally, prediction of RNA binding proteins (RBPs) that induce early splicing changes in C9-ALS reveals signatures consistent with a broad RBP network dysfunction. Together, this work establishes hSpO as an *in vitro* system for investigating early disease mechanisms of ALS in a human multicellular context and highlights dysregulated events that precede cell death.

## Introduction

Amyotrophic lateral sclerosis (ALS) is a fatal neurodegenerative disease characterized by progressive degeneration of upper and lower motor neurons leading to death within 2-5 years of diagnosis (Al-Chalabi & Hardiman, 2013). The most common gene mutation in both familial and sporadic ALS is a hexanucleotide repeat expansion in the *C9orf72* gene, resulting in the production of RNA foci and dipeptide repeat proteins (DPRs) (DeJesus-Hernandez et al., 2011; Renton et al., 2011; van Blitterswijk et al., 2012; Sareen et al., 2013). Several mechanisms have been proposed to explain the progressively degenerative nature of the disease, including disturbances in protein quality control, cytoskeletal defects, excitotoxicity, and dysregulated RNA processing (Kabashi & Durham, 2006; Williamson & Cleveland, 1999; Van Damme et al., 2007; Neumann et al., 2006). Dysregulated RNA processing is exemplified by enrichment of ALS-linked genetic mutations within RNA-binding and RNA-processing genes (Rummens & Da Cruz, 2025; Strong, 2010) and mislocalization and aggregation of the RNA-binding protein (RBP) TDP-43, prevalent in over 97% of all ALS cases (Neumann et al., 2006; Barmada & Finkbeiner, 2010; Polymenidou et al., 2011). However, the study of these disease mechanisms has largely been limited by the lack of a model that allows for the interrogation of longitudinal changes in a human multicellular context.

We previously developed spinal cord/hindbrain organoids (hSpO), a human induced pluripotent stem cell (hiPSC)-derived multicellular model that recapitulates the cell diversity present in the human spinal cord, including cholinergic motor neurons, as well as glutamatergic, and GABAergic interneurons (Andersen et al., 2020). The ability to culture these multicellular organoids long-term unlocks previously inaccessible timelines that allow us to track cellular and molecular changes over time, and potentially highlight early-stage disease mechanisms.

Here, we use hSpO derived from healthy control and *C9orf72* ALS (C9-ALS) hiPSCs to investigate a longitudinal trajectory of C9-ALS. First, we validate the presence of key *C9orf72*-related pathologies including RNA foci, DPRs, cell death, and dysfunction of neuronal processes. Next, by transcriptionally profiling hSpO over time, we identify a progression of disease processes, with RNA splicing emerging as an early dysregulated event. Finally, we show that prediction of RNA binding protein (RBP) binding to dysregulated RNAs reveals signatures consistent with a broad RBP network dysfunction that precedes cell death. Altogether, this work establishes a human multicellular model of C9-ALS and highlights early molecular events preceding cell death.

## Results

### Spinal cord organoids derived from C9-ALS hiPSCs show key *C9orf72*-related pathologies

To establish a human, multicellular model of *C9orf72* disease progression, we generated hSpO as previously described (Andersen et al., 2020; Pali et al., 2026; Peters et al., 2025; Pio et al., 2026) (Figure 1A) and investigated *C9orf72*-related pathologies over time. Spinal cord organoids consist of a diversity of neurons, including cholinergic motor neurons and glutamatergic and GABAergic interneurons, shown by immunohistochemistry for NF-H and MAP2 over-layed with CHAT, GABA, and VGLUT2 in day 61 hSpO cryosections (Figure 1B). We generated hSpO derived from six healthy control and six C9-ALS hiPSC lines (Figure 1A) and validated correct patterning into spinal cord fate by performing bulk RNA sequencing at days 30, 50, and 75 in vitro (at least 2 differentiation batches were collected per hiPSC line, with 2-4 biological replicates per line and batch; Table S1A, B lists hiPSC lines used in various experiments, and Table S1C details samples used for bulk RNA sequencing). Principal component analysis (PCA) demonstrated separation of samples by organoid age (Figures 1C and S1A, B). Normalized expression of select neuronal genes over time, including the synaptic marker *SNAP25*, the vesicular acetylcholine transporter *SLC18A3*, the vesicular glutamate transporter *SLC17A6*, and the vesicular GABA and glycine transporter *SLC32A1*, demonstrated the presence of expected neuronal populations in both control and C9-ALS hSpO, with comparable expression patterns between conditions (Figure S1C).

**Figure 1:**
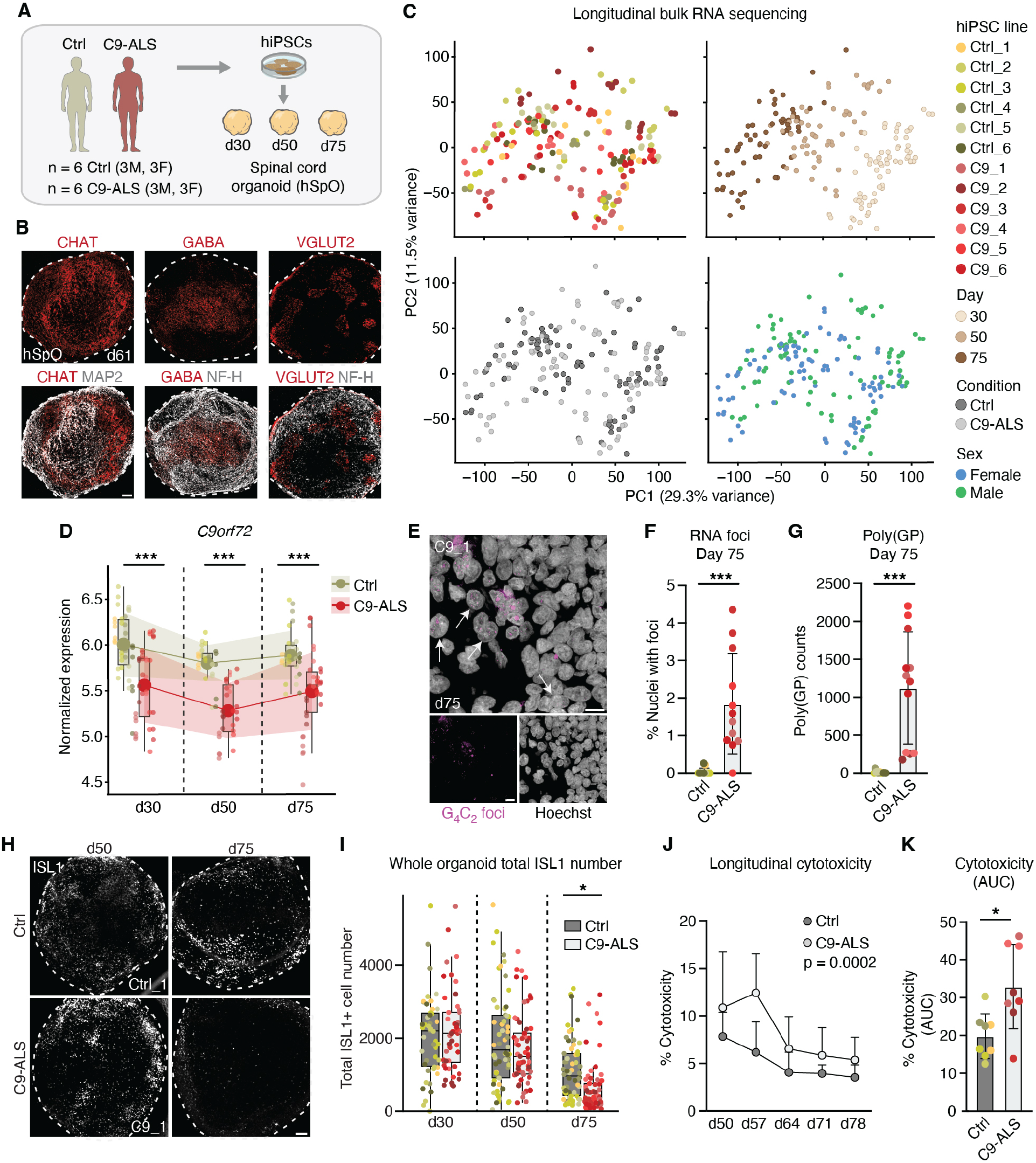
C9-ALS hSpO recapitulate *C9orf72*-related pathologies and cell death. **(A)** Schematic of Ctrl and C9-ALS hiPSC lines used to generate human spinal cord organoids (hSpO). **Table S1A** lists hiPSC lines used in various experiments. **(B)** Representative immunohistochemistry images of cholinergic (CHAT), GABAergic (GABA) and glutamatergic (VGLUT2) neurons (MAP2, NF-H) in hSpO cryosections at day 61. **(C)** Principal Component Analysis (PCA) plots from bulk RNA sequencing showing control and C9-ALS hSpO colored by either hiPSC line they were derived from, age, sex, or condition. Each dot in the PCA represents a sample (n = 95 C9-ALS samples and n = 92 control samples from 6 control and 6 C9-ALS lines, with 1-2 differentiation batches and 2-4 replicates per line, batch and timepoint). Each sample represents 2-3 organoids pooled together. **(D)** Boxplot showing trimmed mean of M-values (TMM) normalized expression values for *C9orf72* over time in control and C9-ALS hSpO. Each dot represents a sample from **Figure 1C**, colored by the hiPSC line they were derived from. Middle black lines represent median and colored large dots represent mean expression in control (green) and C9-ALS (red) hSpO. Shaded green and red regions represents standard deviation in control and C9-ALS conditions, respectively. **(E)** Representative image of fluorescent in-situ hybridization (FISH) of G_4_C_2_ sense RNA foci overlayed with Hoechst in a C9_1 hSpO cryosection at day 75. **(F)** Bar plot showing the percentage of nuclei with G_4_C_2_+ foci in control and C9-ALS hSpO at day 75. Each dot represents an average number of foci per organoid, with 3 cryosections and one field of view per section counted per organoid. n = 3 organoids per line, with n = 4 hiPSC lines in each control and C9-ALS condition. Wilcoxon matched-pairs signed rank test (***p = 0.001). **(G)** Bar plot showing immunoassay poly(GP) counts in control and C9-ALS hSpO at day 75. Each dot represents a replicate per line. n = 2 biological replicates per hiPSC, with n = 4 hiPSC lines in each control and C9-ALS condition. Each replicate represents 2 organoids pooled together. Wilcoxon matched-pairs signed rank test (***p = 0.0005). **(H)** Representative images showing immunohistochemistry for ISL1 in whole control and C9-ALS hSpO at days 50 and 75. **(I)** Boxplot showing counts of total ISL1 numbers in whole hSpO over time. Each dot represents one organoid. n = 166 control and n = 166 C9-ALS organoids across all time points in 2-4 batches. n = 6 hiPSC lines in each control and C9-ALS condition. Two-way ANOVA with Day (F(2, 326) = 43.19, ***p < 0.001) and Condition (control vs. C9-ALS; F(1, 326) = 3.35, p = 0.07) as factors (Day × Condition interaction, F(2, 326) = 2.09, p = 0.13). Pairwise comparisons of estimated marginal means (control vs. C9-ALS) within each timepoint were performed with Tukey adjustment for multiple comparisons (day 30: p = 0.58, day 50: p = 0.26, day 75: *p = 0.016). **(J)** Line graph showing percent cytotoxicity in control and C9-ALS hSpO over time measured by lactate dehydrogenase (LDH) assay. Each dot represents an average of n = 4 control and n = 4 C9-ALS hiPSC lines, each with 2 replicates. Each replicate was collected from a well of a 24-well plate with x2 hSpO each. A mixed-effects analysis was performed with Day (F(1.99, 59.68) = 27.22, ****p < 0.0001) and Condition (control vs. C9-ALS; F(1, 30) = 17.43, ***p = 0.0002) as factors (Day × Condition interaction, F(1.99, 59.68) = 3.76, *p = 0.029), followed by Tukey’s multiple comparisons test per each timepoint (day 50: p = 0.07; day 57: ****p < 0.0001; day 64: *p = 0.02; day 71: *p = 0.02; day 78: *p = 0.01). **(K)** Bar plot showing the area under the curve (AUC) of percent cytotoxicity over time. Each dot represents one replicate in the LDH assay. Each replicate was collected from a well with x2 hSpO each. n = 2 replicates per line, with n = 4 hiPSC lines in each control and C9-ALS conditions. Mann-Whitney test (*p = 0.02). Scale bars = 10 µm **(E)**, 100 µm **(B, H)**

We next validated the presence of key *C9orf72*-relevant pathologies. Southern blot analysis in hiPSC lines confirmed the presence of the hexanucleotide repeat expansion that was maintained after differentiation into hSpO (Figure S1D). Consistent with this finding, expression of *C9orf72* was found to be significantly downregulated at all timepoints in C9-ALS hSpO compared to healthy controls (Figure 1D). Next, we performed fluorescence in-situ hybridization (FISH) for G_4_C_2_ RNA at day 75 and confirmed that nuclei containing G_4_C_2_ foci were present exclusively in C9-ALS hSpO (Figures 1E, F and S1E), with a range of 2-9 foci per cell (Figures S1F, G). Finally, using immunoassay, we confirmed the presence of the poly(GP) dipeptide repeat protein in C9-ALS hSpO at day 75 (Figures 1G and S1H).

Differential gene expression analysis per timepoint confirmed changes consistent with C9-ALS pathology and neuronal dysfunction (Figure S2A and Tables S1D–F). For example, *CHCHD2*, a gene important for mitochondrial respiration in motor neurons and associated with a pathogenic variant linked to altered calcium homeostasis and TDP-43 aggregation, was consistently downregulated in C9-ALS hSpO across all timepoints (Harjuhaahto et al., 2020; Ikeda et al., 2024) (day 30: ***adj. p-val = 0.0009, day 50: *adj. p-val = 0.046, day 75: **adj. p-val = 0.008). *TIA1*, an ALS-associated RNA binding protein found to be sequestered by TDP-43’s 35-kDa truncated fragment into cytoplasmic inclusions (Jiang et al., 2022; Mackenzie et al., 2017), was upregulated at day 30 in C9-ALS hSpO (day 30: *adj. p-val = 0.01, day 50: adj. p-val = 0.82, day 75: adj. p-val = 0.97), while the TDP-43 target *UNC13B*, was downregulated at day 50 (Brown et al., 2022; O’Neill et al., 2025) (day 30: adj. p-val = 0.77, day 50: *adj. p-val = 0.014, day 75: adj. p-val = 0.151). *DNAJC7*, a member of the heat shock protein family identified as a risk gene for ALS, was upregulated at day 75 in C9-ALS hSpO (Farhan et al., 2019) (day 30: adj. p-val = 0.11, day 50: adj. p-val = 0.53, day 75: *adj. p-val = 0.03). At this timepoint, we also observed a significant reduction in *NEFL* (day 30: adj. p-val = 0.30, day 50: adj. p-val = 0.25, day 75: *adj. p-val = 0.045), and a trend towards downregulation of other neuronal and motor neuron markers including *ISL1* (day 30: adj. p-val = 0.59, day 50: adj. p-val = 0.09, day 75: adj. p-val = 0.09) and *SLC5A7* (day 30: adj. p-val = 0.59, day 50: adj. p-val = 0.10, day 75: adj. p-val = 0.09), suggesting a shift in the motor neuron population in C9-ALS over time.

To explore this further we performed immunohistochemistry for the motor neuron marker ISL1 in whole hSpO and quantified total numbers over time. We found that ISL1^+^ cell numbers were comparable in control and C9-ALS hSpO at days 30 and 50, but were significantly decreased in C9-ALS at day 75 (*p-val = 0.016; Figures 1H, I and S2B). Finally, we assessed cell death by measuring lactate dehydrogenase (LDH) release into the culture media and found an overall increase in C9-ALS hSpO (Figures 1J, K and S2C), with a peak at day 57 (**p-val = 0.006; Figure S2D), preceding the reduction in ISL1^+^ cell numbers. Overall, our analysis indicates that C9-ALS hSpO display *C9orf72*-related pathologies including presence of RNA foci and DPRs, as well as longitudinal changes suggesting neuronal dysregulation and cell death over time.

### Distinct time-dependent C9-ALS gene modules identified by longitudinal network analysis

To investigate molecular and cellular processes associated with longitudinal changes in C9-ALS hSpO, we next generated a gene co-expression network using all bulk RNA sequencing samples at days 30, 50 and 75. Weighted gene co-expression analysis (WGCNA) allows us to identify modules of co-expressed genes that are correlated with age in C9-ALS organoids. With this analysis, we identified 40 distinct gene modules with a range from 3,054 genes in M1 to 125 genes in M40 (Figures 2A and S3A; the full list of genes per module and their correlation values are shown in Table S1G). Modules were grouped into three main clusters based on eigengene hierarchical clustering, with eigengenes being the first principal component of all genes in a given module (Figure 2A). Overall, we found 8 modules that showed a significant correlation with disease state irrespective of time, and 10 modules that showed a significant correlation at a specific timepoint with C9-ALS (Figure 2A), suggesting that some processes may be dysregulated at distinct points in time in the disease trajectory.

**Figure 2:**
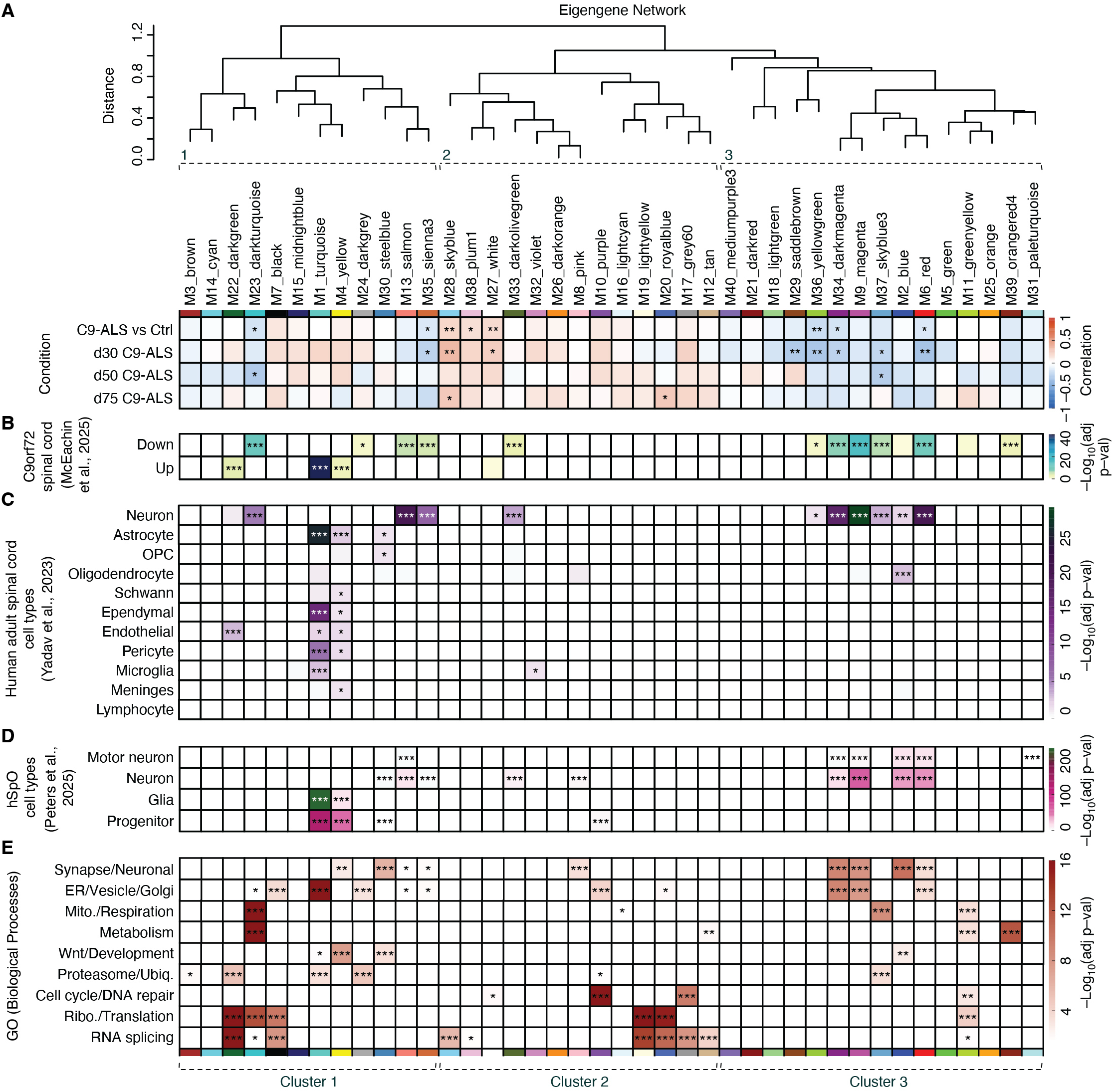
Gene co-expression network in C9-ALS hSpO over time. **(A)** Cluster dendrogram illustrating the similarity of network modules from WGCNA based on correlations between eigengenes, the first principal component of each module. Clusters of related modules are indicated by dashed lines. Correlations between traits of disease state (C9-ALS vs control), and age (d30 C9-ALS, d50 C9-ALS, d75 C9-ALS) were assessed using biweight midcorrelation (BiCor) analysis and represented as a heatmap. Heatmap shows BiCor correlation coefficient between module eigengenes and traits. P-values indicate significance of each module-trait relationship (*p < 0.05, **p < 0.01). **(B)** Heatmap showing Fisher’s exact test (FET) enrichment of proteins that were either increased or decreased in C9orf72 patient spinal cords (McEachin et al., 2025) within network modules. Significance was determined by one-tailed FET, and heatmap is colored based on -log_10_(adj. p-value). **(C)** Heatmap showing FET enrichment of top marker genes from adult spinal cord cell types (Yadav et al., 2023) within network modules. Significance was determined by one-tailed FET, and heatmap is colored based on -log_10_(adj. p-value). **(D)** Heatmap showing FET enrichment of top marker genes from day 60-80 hSpO cell types (Peters et al., 2025) within network modules. Significance was determined by one-tailed FET, and heatmap is colored based on -log_10_(adj. p-value). **(E)** Heatmap showing gene ontology (biological process) enrichment terms for modules identified by WGCNA. Enriched terms were grouped into broader functional categories (rows) based on keyword matching to GO term descriptions (e.g., “Ribosome/Translation”, “Synapse/Neuronal”; full results of GO terms for all modules in **Table S1K**). For modules with multiple significant terms within a category, the term with the lowest adjusted p-value is shown. Statistical significance was assessed via Bonferroni-adjusted p-values (α = 0.05 cutoff for term inclusion) and heatmap is colored based on -log_10_(adj. p-value), (*p < 0.05, **p < 0.01, ***p < 0.001).

To characterize the modules further, we performed Fisher’s exact test (FET) for (i) proteins found to be up- or down-regulated in *C9orf72* postmortem spinal cord (McEachin et al., 2025) (Figure 2B and Table S1H), (ii) top marker genes in human adult spinal cord cell types (Yadav et al., 2023) (Figure 2C and Table S1I), (iii) top markers in hSpO cell types from single cell RNA sequencing (Peters et al., 2025) (Figure 2D and Table S1J), and (iv) biological processes from gene ontology (GO) analysis (Figure 2E and Table S1K). Cluster 1 showed modules with enrichment for glial markers; as well as modules with enrichment for neuronal markers, many of which also showed overlap with proteins downregulated in the *C9orf72* human postmortem spinal cord (Figure 2C, D), including *SV2A, THY1*, and *SYNGR1* in M35-sienna3. Similarly, modules in cluster 3 were enriched for neuronal genes that were downregulated in the postmortem spinal cord, as well as neuron-specific genes from hSpO, and included *SNAP91, CAMK2B*, and *SYP* in M34-darkmagenta, and *ACHE* and *DLG4* in M6-red. In contrast, modules in clusters 1 and 3 showed limited overlap with proteins upregulated in postmortem spinal cord, signatures dominated by glial reactivity and neuroinflammation that are known to be highly upregulated in ALS spinal cord (Humphrey et al., 2023; Tam et al., 2019; O’Neill et al., 2025; Zelic et al., 2025). This is consistent with the relative absence of glial cell types and glial reactivity in hSpO at the timepoints assayed (Andersen et al., 2020; Peters et al., 2025; Pio et al., 2026). Lastly, modules in cluster 2 did not show abundant significant cell type enrichment, suggesting representation of more ubiquitously expressed genes. Additionally, the limited enrichment for postmortem spinal cord signatures in this cluster more broadly suggests that these modules may identify early-stage disease processes not yet captured in postmortem tissue, typically representing end-stage pathology. Overall, this analysis identifies modules representing cell- and stage-specific co-expressed genes that show concordance with *C9orf72* patient tissue.

### Modules correlated with C9-ALS show progressive dysregulation of synaptic transmission, cellular respiration, and RNA metabolism programs

To more comprehensively explore modules associated with C9-ALS pathology and define their longitudinal patterns, we plotted eigengene levels over time for all modules and focused on those that showed a significant difference between control and C9-ALS in at least one timepoint (Figures 3A, B and S4A, B). Overall, we observed neuron-enriched modules in clusters 1 and 3 to be downregulated in C9-ALS (Figures 3A and S4A). Conversely, ubiquitously expressed modules in cluster 2 were upregulated in C9-ALS (Figures 3B and S4B). To understand what processes were represented in these modules over time, we next performed gene ontology enrichment. At day 30, we observed downregulated processes in C9-ALS that were related to protein localization, synapse organization, and ion transport (M35-sienna3, M34-darkmagenta, M6-red), alongside modules reflecting energy metabolism, mitochondrial electron transport chain, and oxidative phosphorylation (M36-yellowgreen, M23-darkturquoise, M37-skyblue3). C9-ALS upregulated processes at day 30 were linked to DNA repair mechanisms (M27-white), as well as to spliceosome function and RNA splicing (M28-skyblue), highlighting early RNA splicing dysfunction in C9-ALS hSpO (Figure 3C). At day 50, C9-ALS module eigengenes were associated with downregulation of electron transport chain and ATP synthesis pathways, highlighting a persistence of mitochondrial dysfunction (M23-darkturquoise, M37-skyblue3; Figure 3C). Finally, at day 75 upregulated modules were related to the assembly of RNA and protein complexes, including ribosome biogenesis (M20-royalblue), indicating that RNA processing dysregulation persists into later stages of C9-ALS hSpO.

**Figure 3:**
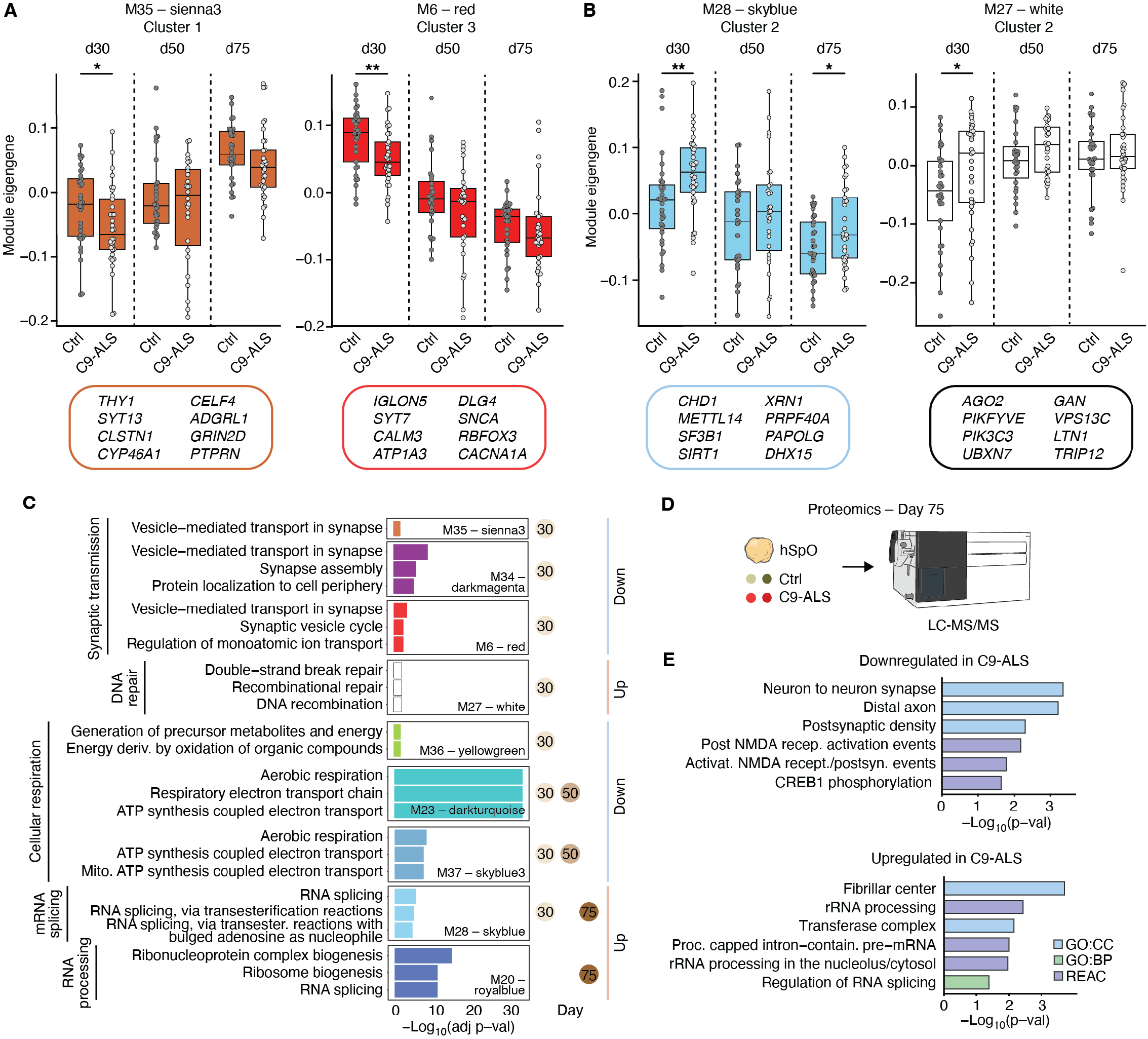
Gene co-expression network analysis reveals progression of disease processes in C9-ALS hSpO. **(A)** Boxplots showing module eigengenes that are decreased in C9-ALS hSpO from clusters 1 and 3. Boxes represent the median and interquartile range (IQR: 25th-75th percentile), with whiskers extending to 1.5 × IQR. Example genes per module are shown in corresponding colored boxes below. Each dot represents a sample from **Figure 1C**. T-tests were performed for each timepoint independently (*p < 0.05, **p < 0.01). Boxplots for other modules in these clusters are shown in **Figure S4A. (B)** Boxplots showing module eigengene that are increased in C9-ALS hSpO for two representative modules from cluster 2. Boxes represent the median and interquartile range (IQR: 25th-75th percentile), with whiskers extending to 1.5 × IQR. Example genes per module are shown in corresponding colored boxes below. Each dot represents a sample from **Figure 1C**. T-tests were performed for each timepoint independently (*p < 0.05, **p < 0.01). Boxplot for the other module in this cluster is shown in **Figure S4B. (C)** Bar plots showing gene ontology (biological process) terms for modules that show differential eigengene levels between control and C9-ALS in at least one timepoint. Colored circles on the right indicate the timepoint(s) at which each module shows significant differential eigengene expression, and vertical colored lines indicate the direction of change (red = upregulated, blue = downregulated). **(D)** Schematic showing design for proteomics experiment on day 75 hSpO. n = 3 replicates per hiPSC line, with n = 2 hiPSC lines in each control and C9-ALS condition. Each replicate represents 2-3 organoids pooled. **(E)** Gene Ontology terms for proteins downregulated and upregulated in C9-ALS, colored by reference source used.

To validate some of the processes observed at the gene network analysis, we performed small-scale proteomics at day 75 in healthy control and C9-ALS hSpO (Figure 3D and Table S1L–N). We found processes identified by the network analysis also dysregulated at the protein level at this timepoint. Neuronal-associated terms, such as activation of NMDA receptors and synaptic transmission-related processes were downregulated (Figures 3E, S4C and Table S1O), while RNA metabolism processes were upregulated in C9-ALS, recapitulatingchanges at the RNA level (Figures 3E, S4C and Table S1P). Altogether, our longitudinal systems-level analysis of bulk transcriptomics and proteomics in C9-ALS hSpO, highlighted a temporal progression of disrupted cellular and molecular events preceding neuronal dysfunction and cell death. Notably, RNA splicing dysregulation emerged as an early and persistent process throughout disease progression.

### RNA splicing dysregulation in C9-ALS hSpO recapitulates splicing changes found in *C9orf72* tissue

Prominent RNA metabolism dysregulation found in our network analysis prompted us to investigate TDP-43 pathology in hSpO. TDP-43 is an RNA-binding protein that mislocalizes from the nucleus to the cytoplasm in a majority of ALS cases, including C9-ALS (Al-Sarraj et al., 2011; Murray et al., 2011; Neumann et al., 2006). To investigate the subcellular localization of TDP-43 in C9-ALS hSpO we first performed fractionation of nuclear and cytoplasmic components followed by Western blot for TDP-43. We observed a significant increase in cytoplasmic TDP-43 across all timepoints in C9-ALS, along with a reduction in the nuclear/cytoplasmic ratio (Figures 4A and S5A). Notably, we observed no significant change in nuclear TDP-43 levels over time (Figure S5B). However, recent analysis of postmortem tissue indicates that overt loss of nuclear TDP-43 occurs in a small fraction of cells (Liu et al., 2019;

**Figure 4:**
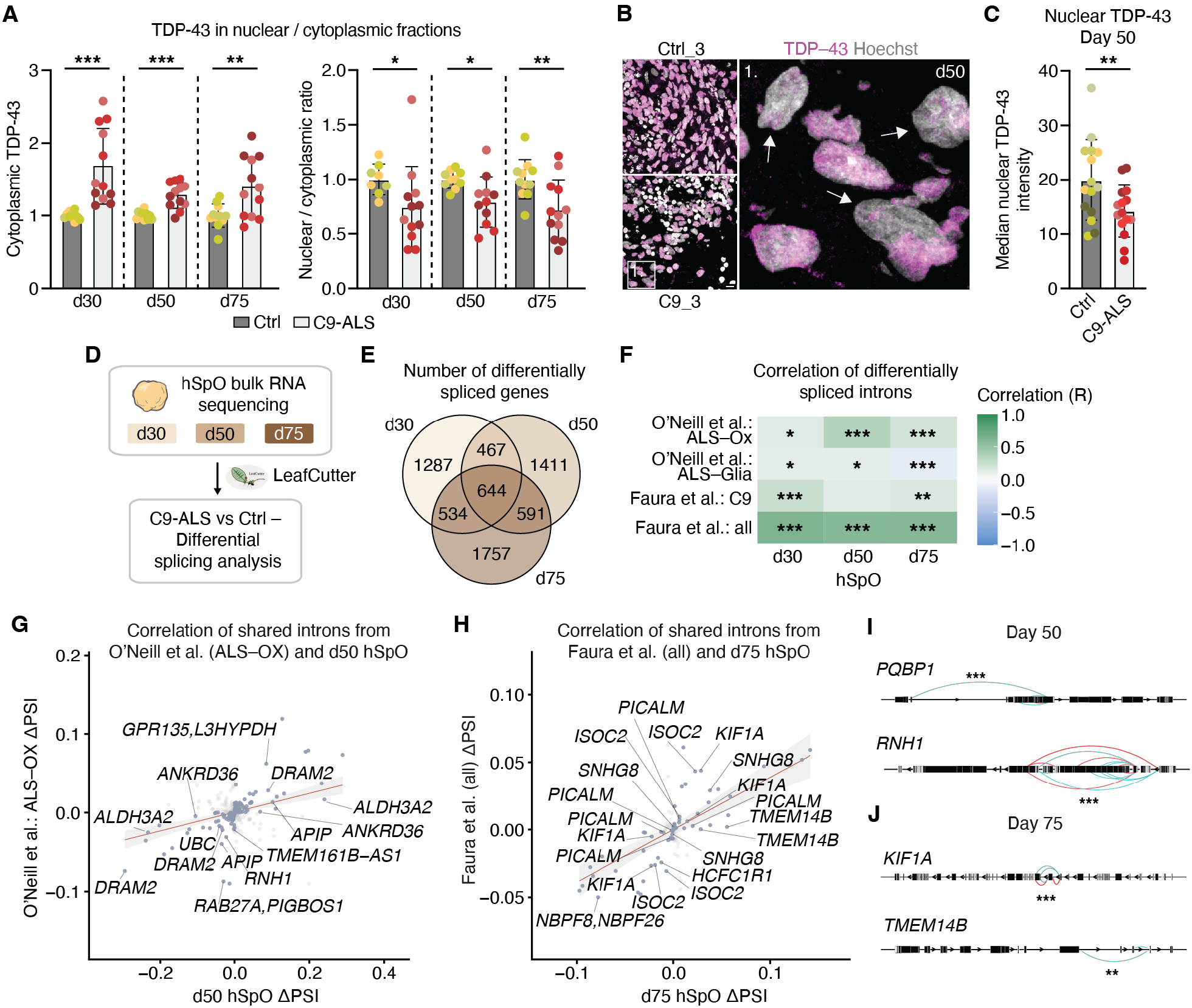
C9-ALS demonstrates TDP-43 mislocalization and dysregulated RNA splicing. **(A)** Bar plots showing quantification of cytoplasmic TDP-43 (left) and nuclear/cytoplasmic TDP-43 ratio (right) from TDP-43 Western blots in cytoplasmic and nuclear fractions. Each dot represents a replicate, with 5 organoids pooled and fractionated per replicate. n = 2-4 replicates per hiPSC line, with n = 3 hiPSC lines in each control and C9-ALS condition. All C9-ALS samples were normalized to batch controls per timepoint. Quantification of nuclear TDP-43 is shown in **Figure S5B**. For cytoplasmic TDP-43, a mixed effects analysis with Day (F(1.44, 43.89) = 2.56, p = 0.104) and Condition (control vs C9-ALS; F(1, 61) = 36.96, ****p < 0.0001) as factors was performed (Day × Condition interaction, F(1.44, 43.89) = 2.56, p = 0.104) followed by Tukey’s multiple comparisons test at each timepoint: day 30, ***p = 0.0009; day 50, ***p = 0.0002; day 75, **p = 0.0087. For the nuclear/cytoplasmic TDP-43 ratio, a mixed-effects for Day (F(1.524, 28.19) = 0.1180, p = 0.8348), and Condition (control vs C9-ALS; F(1, 21) = 15.69, ***p = 0.0007) as factors was performed (Day × Condition interaction, F(1.524, 28.19) = 0.12, p = 0.84) followed by a Tukey’s multiple comparisons test at each timepoint: day 30, *p = 0.049; day 50, *p = 0.016; day 75, **p = 0.009. **(B)** Representative images showing immunohistochemistry for TDP-43 and Hoechst in day 50 control and C9-ALS hSpO cryosections. Inset shows examples of reduced TDP-43 nuclear intensity in C9-ALS hSpO. **(C)** Bar plot showing median nuclear intensity of TDP-43 in day 50 control and C9-ALS hSpO. Each dot represents the median intensity value per organoid, with 3 organoids per hiPSC line, and n = 5 hiPSC lines in each control and C9-ALS condition. For each organoid, three cryosections were imaged and quantified. Wilcoxon matched-pairs signed rank test (**p < 0.01). **(D)** Schematic of differential splicing analysis pipeline performed on all samples from **Figure 1C** using Leafcutter. **(E)** Venn diagram showing the number of differentially spliced genes over time from Leafcutter. Significantly spliced introns were filtered (adj. p-val < 0.05) and their corresponding genes counted. **(F)** Heatmap showing the correlation of top hSpO differentially spliced introns (adj. p-val < 0.01) over time with human postmortem brain and spinal cord datasets. Pearson’s correlation coefficient (R) and p-values were calculated for each timepoint (*p < 0.05, **p < 0.01,***p < 0.001). Number of introns per dataset are as follows: n = 34–96 (Faura et al., “all”), n = 198–423 (Faura et al., C9 carriers) n = 530–966 (O’Neill et al., ALS-Glia), n = 272–602 (O’Neill et al., ALS-Ox), with range indicating timepoints. **(G)** Scatterplot showing correlation of day 50 hSpO and ALS-Ox (O’Neill et al., 2025) differentially spliced introns, using Δ percent spliced in (ΔPSI) for each. Each point represents an individual splicing event, with positively correlated introns highlighted in dark gray, line of best fit shown in red, and corresponding gene names indicated. Pearson’s correlation coefficient R = 0.35, p = 3.18 × 10^-9^, n = 272 splicing events. **(H)** Scatterplot showing correlation of day 75 hSpO and Faura et al., “all” (2025) differentially spliced introns, using Δ percent spliced in (ΔPSI) for each. Each point represents an individual splicing event, with positively correlated introns highlighted in dark gray, line of best fit shown in red, and corresponding gene names indicated. Pearson’s correlation coefficient R = 0.59, p = 3.8 × 10^-10^, n = 96 splicing events. **(I)** Examples of differentially spliced introns from day 50 hSpO C9-ALS shown for *PQBP1* and *RNH1*. Colored lines represent an increased intron inclusion in C9-ALS (red) or a decreased intron inclusion in C9-ALS (blue). Significance per splice junction from Leafcutter is denoted (*** adj. p-value < 0.001). **(J)** Examples of differentially spliced introns from day 75 hSpO C9-ALS shown for *KIF1A* and *TMEM14B*. Colored lines represent an increased intron inclusion in C9-ALS (red) or a decreased intron inclusion in C9-ALS (blue). Significance per splice junction from Leafcutter is denoted (** adj. p-value < 0.01, *** adj. p-value < 0.001). Scale bar = 10 µm **(B)**

Rothstein et al., 2025), and might be masked by bulk protein analysis. Therefore, we next explored nuclear TDP-43 at an individual cell level by performing immunohistochemistry of control and C9-ALS hSpO cryosections at day 50. We measured TDP-43 nuclear fluorescence intensity and observed a decrease in median nuclear TDP-43 in C9-ALS hSpO (Figures 4B, C and S5C).

Next, we investigated whether this partial nuclear TDP-43 loss contributes to an increased inclusion of cryptic exons, as previously shown (Baughn et al., 2023; Bryce-Smith et al., 2025; Ma et al., 2022; Mehta et al., 2023). Quantitative RT-PCR of known cryptic events in *STMN2* and *ELAVL3* over time showed a significant increase in C9-ALS hSpO at day 75, however this upregulation was primarily driven by one hiPSC line (Figure S5C). In addition, other known cryptic events such as *KALRN* showed no significant increase (Figure S5C). We reasoned that the RNA splicing dysregulation signatures observed above in C9-ALS hSpO might be driven by less canonical splicing changes, and therefore used the R package Leafcutter (Y. I. Li et al., 2018) to unbiasedly detect transcriptome-wide variation in splicing. Using our bulk RNA sequencing dataset, we compared all six C9-ALS lines to six control lines at days 30, 50, and 75 (Figure 4D and Table S1Q–S). Over 600 genes showed significant differential intron splicing shared across all timepoints, with an overall increase in the number of differentially spliced genes over time (Figures 4E and S6A). Genes with differentially spliced introns that were common to all timepoints were enriched in processes related to cilia, axonogenesis, and neuronal projections (Figure S6B). On the other hand, genes related to protein localization showed greater differential splicing at early timepoints, while nucleotide excision repair, and actin polymerization genes were more significantly spliced at later timepoints (Figure S6B). Correlation of the top differentially spliced introns in hSpO (adj. p-val < 0.01) with those found in postmortem ALS tissue indicated that splicing dysregulation in hSpO was overall concordant to that seen in patient tissue (Figure 4F). We saw the highest correlation with splicing events found in the frontal cortex of FTLD-TDP patients from Faura et al., (2025), followed by events in the spinal cord of the ALS-Ox subtype described in O’Neill et al., (2025). Splicing events found in *C9orf72* expansion carriers from Faura et al., (2025) also showed a significant correlation with hSpO at days 30 and 75. Exploration of shared differentially spliced introns between hSpO and patient datasets highlighted the presence of splicing events in ALS-relevant genes, including the motor protein *KIF1A* (Zheng et al., 2024), as well as the primate-specific transmembrane protein 14B (*TMEM14B*), and genes involved in RNA processing, including *PQBP1* and *RNH1* (Figures 4G–J).

Having established the *in vivo* relevance of differentially spliced intron events in hSpO, we next investigated the prevalence of TDP-43-relevant events. We compared significant events in hSpO at days 30, 50 and 75, with alternatively spliced genes found in FACS-sorted TDP-43 negative neuronal nuclei from FTD-ALS postmortem brain tissue from Ma et al., (2022). Interestingly, despite lower nuclear/cytoplasmic TDP-43 ratios at all timepoints in hSpO, only a small proportion of differentially spliced introns from this postmortem dataset were also significant in hSpO (day 30 = 3.03%, day 50 = 1.51%, day 75 = 0%; Figure S6C). We reasoned that differences between hiPSC lines, as observed in our RT-qPCRs, might be driving the reduced detection of significant events, and therefore next investigated per-line events by performing Leafcutter in individual C9-ALS hiPSC lines compared to all controls per timepoint. Indeed, this analysis revealed differences between hiPSC lines, with some of them showing a higher proportion of TDP-43-related alternatively spliced introns overall (day 30 = 0 – 22.7%, day 50 = 0 – 13.6%, day 75 = 1.5

– 16.7%; Figure S6C). Nevertheless, canonical cryptic events previously described for the spinal cord were not found to be significant in hSpO even in the per-line analysis (e.g. *STMN2* and *UNC13B*) (O’Neill et al., 2025). Altogether, our analysis indicates that hSpO recapitulate certain aspects of RNA processing dysfunction found in ALS brain and spinal cord, including TDP-43 mislocalization and altered intron splicing. Splicing alteration, however, did not fully recapitulate known TDP-43 loss-of-function signatures, which led us to investigate RNA binding proteins more broadly.

### An early network of RNA-binding proteins is predicted to drive splicing dysregulation in C9-ALS

Our network analysis highlighted RNA metabolism as an early dysregulated process in C9-ALS, yet differential splicing analysis with Leafcutter did not show splicing signatures highly concordant with TDP-43 loss-of-function, suggesting that other RNA-binding proteins (RBPs) might be driving this signature. To investigate what RBPs might instead be driving the splicing changes observed in hSpO, we leveraged CircRNAs Interact with Proteins (CRIP), a sequence-based RBP-binding prediction algorithm (Zhang et al., 2019). Although CRIP was originally trained on circular RNA (circRNA)-RBP interactions, its prediction depends only on primary sequence and was therefore applied directly to the splice-site flanking sites on our dataset. Briefly, we selected the top hSpO differentially spliced introns (|ΔPSI| > 0.2, adj. p-val < 0.05) identified in our Leafcutter analysis at each timepoint, and extracted the 400-nucleotide (nt) genomic sequence upstream and downstream of the splice junctions. For each 400-nt flank, a total of 300 overlapping windows were tiled (101-nt sliding window and 1-nt stride), and each window was scored against 137 RBPs using CRIP (Figure 5A). RBP-binding probability was then predicted from RNA linear sequences using the CRIP deep-learning model that scores peak count and binding intensity for both upstream and downstream regions (Figures 5B, S7A and Table ST–V). We used unchanged introns (adj. p-val > 0.09) as background introns for comparison. Finally, to understand the interactions of identified RBPs with each other, we performed STRING network analysis at each timepoint, where RBPs from both CRIP metrics are represented by nodes, and the functional and physical associations of the RBPs to each other are represented by edges (Figure 5C).

**Figure 5:**
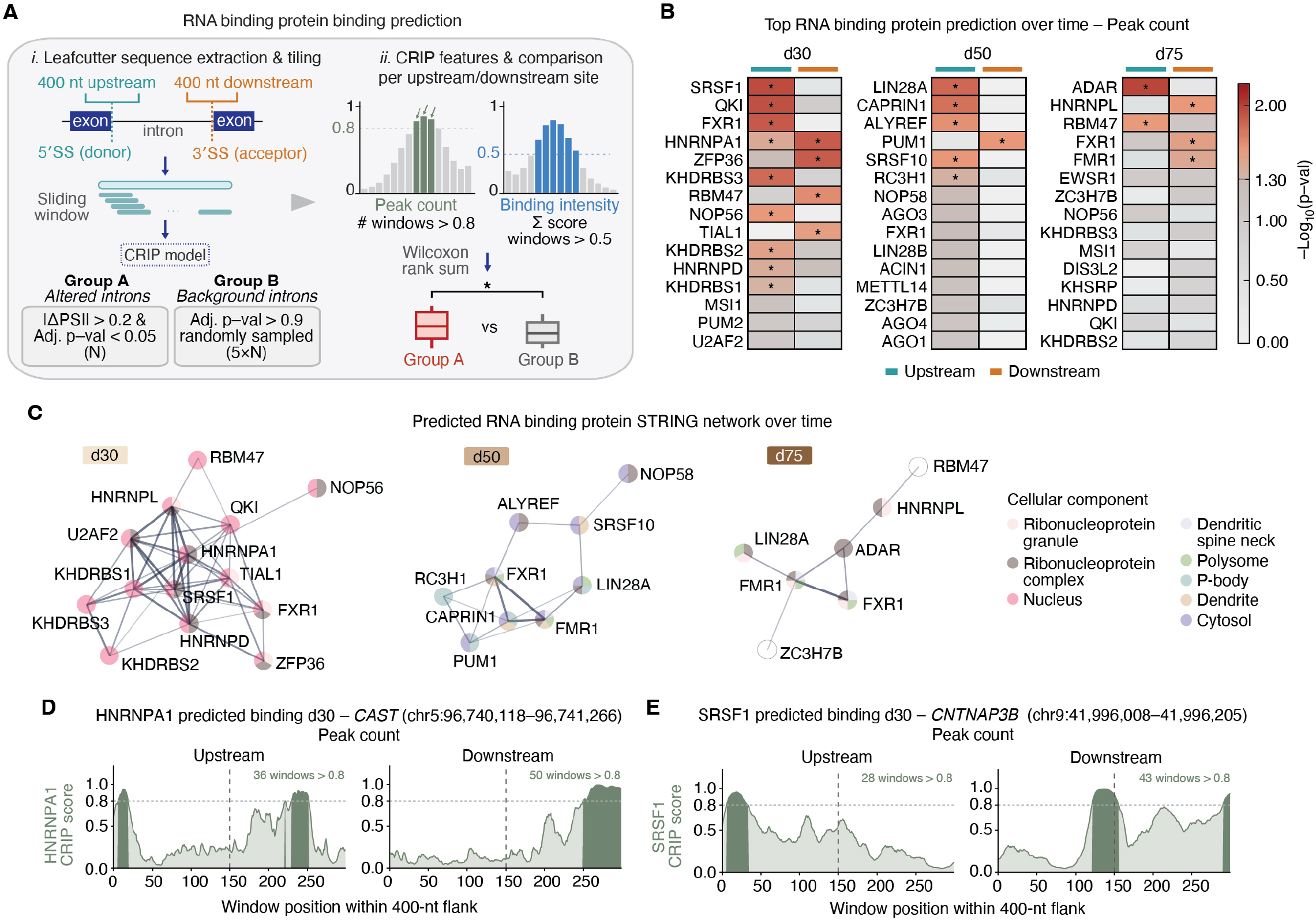
Prediction of RNA binding proteins driving alternative splicing events. **(A)** Schematic illustrating CRIP analysis for the prediction of RNA binding proteins (RBPs) using Leafcutter differentially spliced events. **(B)** Heatmap showing top predicted RBPs per timepoint based on peak counts metric and colored by p-value from Wilcoxon rank sum test comparing peak counts per RBP in altered vs background group of splicing events defined in **A** (*p < 0.05). Dotted line in scale bar legend indicates significance threshold (-log_10_(0.05) = 1.30). Values for both upstream and downstream flanks of splicing event are separately shown. Heatmap of binding intensity metrics shown in **Figure S7A. (C)** STRING network plot showing interactions of RBPs predicted to drive alternative splicing events in hSpO. Significant predicted RBPs from peak counts and binding intensity metrics are shown per timepoint. Each node represents an RBP colored by cellular component gene ontology, and each edge represents a predicted interaction, with thickness denoting strength of data supporting the interaction. **(D)** Line plot showing the predicted binding profile for HNRNPA1 along a *CAST* splicing event site at day 30, shown across 400-nucleotide (nt) upstream and downstream flanks. Per-window CRIP binding probabilities are shown across 300 sliding windows, with darker shading indicating regions where probability score exceed 0.8 (peaks). Vertical dashed lines show the splice junctions per flank. **(E)** Line plot showing the predicted binding profile for SRSF1 along a *CNTNAP3B* splicing event site at day 30, shown across 400-nucleotide (nt) upstream and downstream flanks. Per-window CRIP binding probabilities are shown across 300 sliding windows, with darker shading indicating regions where probability score exceed 0.8 (peaks). Vertical dashed lines show the splice junctions per flank.

This analysis predicted a dynamic range of RBP binding to C9-ALS differentially splicing sites. At day 30, splicing events were strongly associated with the hnRNP family and KH RNA-binding domain-containing RBPs upstream (HNRNPA1, HNRNPD, KHDRBS3, KHDRBS2, KHDRBS1), along with regulators of RNA metabolism such as SRSF1 (Tao et al., 2024) (Figures 5B and S7A). Notably, TARDBP was indeed predicted as one of the top downstream binding RBPs at this timepoint based on binding intensity, although it did not reach significance (upstream p-val = 0.33, downstream p-val = 0.10) (Figure S7A). Day 30 also showed the greatest number of predicted RBPs at any time point, with a network predominately localized to the nucleus and including many ALS-relevant RBPs (Figure 5C). For example, HNRNPA1 and SRSF1 emerged as top hub genes, both demonstrating disease relevance at this stage (Table S1W). Depletion of SRSF1 has been shown to prevent neurodegeneration in both human-derived neurons and drosophila models of *C9orf72* (Hautbergue et al., 2017; Castelli et al., 2021). HNRNPA1 has been found to have ALS-linked mutations (Lee & Rio, 2024) and shown to be modulated by TDP-43 (Deshaies et al., 2018). In our analysis, for example, HNRNPA1 binding was predicted on both upstream and downstream flanks of the calpain inhibitor *CAST* (Figure 5D), whose downregulation contributes to neurodegeneration (Lescouzeres et al., 2025), and SRSF1 binding was predicted on *CNTNAP3B*, a member of the Caspr protein family (Figure 5E), among others.

At day 50, upstream post-transcriptional regulatory RBPs were predicted to drive splice events in hSpO (RC3H1, LIN28A, SRSF10, ALYREF, CAPRIN1, NOP58), with limited downstream RBP signal (PUM1). These RBPs also begin to show p-body, dendritic spine neck, and cytosol involvement, suggesting a shift away from nuclear splice events (Figure 5C). At day 75 we observe an enrichment for neuronal RBPs and polyribosome associating factors downstream (FMR1, FXR1), and ADAR, RBM47 upstream (Figures 5B and S7A). Several RBPs at this later stage are found in ribonucleoprotein granules, defined as membrane-less organelles of RNA and protein, and polysomes, structures of several ribosomes across a single mRNA molecule (Figure 5C). This shift towards translational machinery is further supported by our proteomics data which revealed a concurrent upregulation of rRNA processing at day 75 (Figure 3E).

Overall, our CRIP predictions reveal an early network of RBPs associated with splicing changes prior to detectable neuronal dysfunction. This temporal sequence of predicted RBPs demonstrates a shift from nuclear splicing regulation at day 30, to cytoplasmic RNA processing and translational control later, and warrants investigation of broader RBP dysfunction as an early driver of degeneration in C9-ALS.

## Discussion

Human iPSC-derived multicellular neural organoids have served as powerful models to study neurodevelopment and aspects of neurodegeneration in a human cell-specific context (Birey et al., 2017; Chen et al., 2024; Hruska-Plochan et al., 2024). In C9-ALS, hiPSC-derived models have demonstrated disease-relevant phenotypes such as presence of RNA foci, DPR accumulation and deficits in synaptic function (Szebényi et al., 2021; van der Geest et al., 2024; Gao et al., 2024). However, organoid models of ALS, and related disease frontotemporal dementia (FTD) (Szebényi et al., 2021; van der Geest et al., 2024), have primarily focused on upper motor neurons, and a comprehensive longitudinal analysis of disease signatures that compares to human ALS spinal cord is still lacking. Here, we provide a temporal analysis of multicellular human spinal cord organoids and demonstrate the prevalence of C9-ALS disease hallmarks, neuronal dysfunction and cell death, as well as early and persistent RNA splicing dysregulation predicted to be driven by a dynamic network of RBPs.

Spinal cord organoids ranging from 30-75 days in vitro contain a diversity of neurons that recapitulate the cellular diversity found in the human spinal cord (Andersen et al., 2020; Peters et al., 2025). A significant advantage of 3D cultures is the ability to follow vulnerable cell populations over extended periods of time. Our analysis indicated an overall increase in cell death in C9-ALS organoids and a reduced number of ISL1^+^ cells between days 50 and 75 *in vitro*, suggesting an accelerated decline of motor neurons in C9-ALS. Importantly, our network analysis highlighted several dysregulated processes that preceded cell death in C9-ALS hSpO, many of which have been previously associated with disease. Dysregulation of synapses in C9-ALS was one of the early identified signatures in ALS and has been supported by an overall decrease in synapses, morphological changes, and disturbances in network activity (Henstridge et al., 2018; Bauer et al., 2022; Perkins et al., 2021). Dysregulation of metabolic processes has also been well-established as a contributor to ALS pathogenesis and was recently demonstrated to occur prior to the onset of symptoms (Burg & Van Den Bosch, 2023; Dorst et al., 2023). A prominent signature in our analysis both at early and later timepoints was the presence of RNA metabolism and splicing changes in C9-ALS. RNA metabolism dysregulation is a common hallmark of ALS and overall neurodegeneration (Y. Li & Sun, 2025), with TDP-43 mislocalization and aggregation standing out as central pathologies in a majority of ALS cases and a proportion of FTD and Alzheimer’s disease (AD) (Agra Almeida Quadros et al., 2024; Arai et al., 2006; Neumann et al., 2006). Our data indicates the presence of TDP-43 cytoplasmic mislocalization and a partial reduction in nuclear TDP-43 intensity in C9-ALS hSpO, yet no robust presence of known cryptic events in most hiPSC lines nor TDP-43 aggregation, consistent with results in other 3D cellular models, where TDP-43 dysfunction was either not reported (Bowles et al., 2021; van der Geest et al., 2024) or was induced via injection of patient protein aggregates (Tamaki et al., 2023). Rothstein et al, (Rothstein et al., 2025) recently demonstrated consistent presence of known cryptic exons in *C9orf72* and sporadic hiPSC-derived 2D motor neurons after 46 days *in vitro*. It is possible that a 3D multicellular environment shifts the timeline for cryptic event inclusion in motor neurons and that either longer culture or higher temporal resolution could unmask these events in hSpO. Despite the lack of robust inclusion of canonical cryptic exons, however, our differential splicing analysis revealed a high degree of correlation between splicing changes in hSpO over time and splicing events found in ALS patient tissue, indicating likely relevance to disease. In line with this observation, a large-scale splicing analysis of patient FTLD-TDP frontal cortex indicated that a majority of splicing alterations appeared to be due to general neurodegenerative processes rather than TDP-43 dysfunction (Faura et al., 2025). Notably, our CRIP prediction analysis highlighted RBPs previously linked to ALS, including HNRNPA1, SRSF1, TIAL1 and others, as potential drivers of the observed splice events in hSpO. Indeed, previous analysis of brain patient tissue showed an enrichment of HNRNPH binding motifs in differential splicing events of C9-ALS cases (Prudencio et al., 2015), while RBPs in the SRSF and HNRNP families have previously been shown to be sequestered by G_4_C_2_ repeat RNA in motor neurons (Cooper-Knock et al., 2014). In addition, RBPs identified in our CRIP analysis, including ADAR, HNRNPA1, KHDRBS2, and KHDRBS3, have previously been shown to form complexes with endogenous TDP-43 (Sephton et al., 2011). Building on this, recent findings indicate that mislocalized TDP-43 in the cytoplasm co-sequesters other splicing regulators from the nucleus, including SRSF, HNRNP and TIA RBP family members (Van Zuiden et al., 2025). Early cytoplasmic TDP-43 mislocalization in hSpO is consistent with this observation. Future work will determine whether other RBPs are co-sequestered by TDP-43 in hSpO, and whether such mislocalization contributes to the splicing changes we observe.

Altogether, here we present a temporally resolved analysis of C9-ALS hSpO that reveal early molecular events associated with neurodegeneration in C9-ALS. In our model, an early network of RBPs is predicted to induce alternative splicing changes consistent with those observed in patient tissue. Future work will focus on how hnRNPs, SRSF and TIA networks relate to TDP-43 dysfunction and what sequence of events might precede and directly contribute to neuronal loss.

## Limitations

Organoids derived from hiPSCs follow a mostly developmental timeline. Culture conditions might precipitate degenerative processes, but whether the molecular mechanisms underlying such degenerative processes are comparable to those observed in adult cells *in vivo* is unclear. Organoids lack vasculature, microglia and other immune cells, all of which might play important roles in disease onset and progression *in vivo*. Line-to-line variability introduces an additional limitation that we addressed by incorporating multiple hiPSC lines, batches, and replicates. Finally, the RBP prediction algorithm we employed was trained on a small number of RBPs, limiting the scope of the RBP prediction analysis. In future work, incorporation of isogenic controls and disease-associated mutants will enable more precise, less variable dissection of disease mechanisms contributing to splicing defects.

## Acknowledgements

We would like to thank members of the Andersen and Birey labs for useful discussions and support. We also would especially like to thank Nikolay Tranakiev for support with performing Leafcutter analysis, and Brain Organoid Hub members Alexia King and Zihui Ou for helping generate and maintain organoids used in this study. This work was supported by the Alz-heimer’ Disease Big Data to Biology Training Program (5T32AG087922) to B.A., a grant from the National Institutes of Health (R01AG068247; J.P. and J.J.), and grants from ALS Cure Project (J.A.), The ALS Association (J.A.; doi.org/10.52546/ALSA.24-SGP-690.pc.gr.187896), Edward Mallinckrodt Jr Foundation (J.A.), and Tambourine’s ALS Breakthrough Research Fund in Partnership with the Milken Institute (J.A.).

## Author contributions

B.A., A.M. and J.A. conceived the project and designed the experiments.

B.A. performed experiments and analyzed the data.

S.L. and B.Y. performed and supervised the CRIP analysis.

B.A., A.M., S.S.W. and A.J.R.Y. collected and processed samples for longitudinal bulk RNA sequencing.

S.X.Y., N.E.S., A.J.R.Y., N.E.J.S, and S.H.Z. contributed to the collection, immunohistochemistry, imaging and quantification of ISL1 cells.

S.M. performed the cytotoxicity assay over time.

J.S.T. performed staining, imaging, and quantification of TDP-43 nuclear intensity.

T.P. performed immunohistochemistry for cell type markers.

M.C. and Z.M. performed the DPR assay and RT-qPCRs for cryptic exons and supervised the analysis.

A.N.T. provided guidance on WGCNA and proteomics analysis.

L.Y, C.M.W, and N.T.S. performed the proteomics experiment.

J.P. and J.J. provided support and guidance on FISH.

The B.O.H and S.S.W. generated organoids.

B.A. and J.A. wrote the manuscript with input from all authors.

J.A. supervised the work.

## Competing interest statement

J.A is a named inventor in a patent that covers the generation and assembly of region-specific spinal cord organoids and cortico-spinal assembloids.

## Materials and Methods

### Culture of hiPSCs

Human iPSC lines used in this study were obtained from the Cedars-Sinai Biomanufacturing Center (Table S1A shows information for all lines). All lines were regularly screened and maintained mycoplasma-free. hiPSCs were cultured on Cultrex R&D systems, BME001-05) with daily media changes of mTESR plus (Stemcell Technologies, 100-0276). hiPSC colonies were passaged with ReLeSR (Stemcell Technologies, 05872) upon reaching 80–90% confluency. Detailed protocols available at www.brainorganoidhub.com/protocols.

### Generation of spinal cord organoids (hSpO)

hSpO were generated following previously established protocols (Andersen et al., 2020; Pali et al., 2026; Peters et al., 2025; Pio et al., 2026). Briefly, hiPSC colonies at 80–90% confluency were detached from culture plates using Accutase (BioLegend, 10761-312) and formed into 3D organoids in AggreWell 800 plates (STEMCELL Technologies, Cat. #34815) which is considered as D-1. Next day (D0), the organoids were dislodged into 10 cm plates containing 2.5 µM dorsomorphin (DM, R&D systems, 3093), 10 µM SB-431542 (Selleck Chemicals, S1067) and 10 µM Y-27/Rock Inhibitor (R&D systems, 1254), 1× penicillin/streptomycin (Cytiva, SV30010) in E6 medium (Thermo Scientific, A1516401). From day 1 onwards, Y27 was discontinued. On days 3 and 4, hSpO additionally received 3 µM CHIR-99021 (Selleck Chemicals, S1263). From day 5 to day 9, hSpO received 10 ng/ml FGF2 (R&D systems, 233-FB), 20 ng/ml EGF (R&D systems, 236-EG), 0.1 µM retinoic acid (Sigma Aldrich, R2625), and 3 µM CHIR in NBA+ medium (United States Biological, N1020-02) containing 1× B27 Minus Vitamin A (Thermo Scientific, A3582801), 1× Glutamax (Thermo Scientific, 35050061), 1× penicillin/streptomycin every other day. From day 10 to day 17, hSpO also received 0.1 µM smoothened agonist (SAG, Selleck Chemicals, S7779) every other day. From day 18 to day 23, hSpO received 1× N-2 supplement (Thermo Scientific, 17502-048), 20 ng/ml BDNF (PeproTech, 450-02), 50 µM cAMP (Sigma Aldrich, D0627), 200 µM L-ascorbic acid (L-AA, Sigma Aldrich, A4403), 10 ng/ml IGF-1 (PeproTech, 100-11), and 2.5 µM DAPT (Stemcell Technologies, 100-1046) in NBA+ every other day. From day 24 onwards DAPT was discontinued and hSpO were maintained in NBA media with 1× B27 Minus Vitamin A, 1× Glutamax, 1× penicillin/streptomycin containing BDNF, L-AA, cAMP and IGF-1 at above concentrations.

### Southern blot for *C9orf72* repeat expansion analysis

Genomic DNA was extracted from hSpO using a DNeasy Blood and Tissue Kit (Qiagen, 69504) and outsourced to Celplor LLC for Southern blot analysis. Probe labeling of (GGGGCC)5 repeats, Southern blotting and signal recording were performed following Celplor established protocols.

### RNA collection, extraction and bulk RNA sequencing

Organoids were placed in 1.7 mL tubes at designated timepoints (days 30, 50, 75), all media was removed, and samples were snap frozen on dry ice and stored at -80 °C. Total RNA was isolated using the RNeasy Mini Kit and RNase-Free DNase Set (Qiagen, 74106) and stored at -80°C until sequencing. RNA was shipped to Admera Health for library preparation and sequencing. RNA integrity was assessed and samples with RIN values greater than or equal to 8 were used for library preparation. Messenger RNA was enriched via poly(A) selection, and strand-specific libraries were prepared using the NEBNext Ultra II Directional RNA Library Prep Kit. Libraries were sequenced on an Illumina platform using paired-end 2×150 bp chemistry, targeting a sequencing depth of 40 million paired end reads per sample.

### Quantitative PCR for cryptic exons

Template cDNA was prepared by reverse transcription using the Maxima First Strand cDNA Synthesis Kit (Life Technologies, K1672) with 200 ng of RNA and stored at -20°C. Expression of mRNA for cryptic exons was quantified by quantitative PCR (qPCR) using custom TaqMan gene expression assays (Thermo Fisher Scientific) on a QuantStudio 6 Flex system (Applied Biosystems). For each well, 2 µL of cDNA was loaded. Custom TaqMan sequences for truncated-*STMN2* (APZTKFJ), *ELAVL3-CE* (AP7DWP9), and *KALRN-CE* (APKCDUH) are: truncated-*STMN2* Forward: 5’-TGCACATCCCTACAATGGCTAAAA-3’; truncated-*STMN2* Reverse: 5’-TCTACCTTTCTCTCGAAGGTCTTCTG-3’; truncated-*STMN2* reporter probe: 5’-CCGAGTCCCATTGCTG-3’; *ELAVL3-CE* Forward: 5’-GCCTCAAATTACAGACGAAGACCAT-3’; *ELAVL3-CE* Reverse: 5’-GGTCACAGCCGGAGTCA-3’; *ELAVL3-CE* Reporter probe: 5’-CAGTGTCACATGCACCTTG-3’; *KALRN-CE* Forward: 5’-CGGAGTTTGTGCGACTTCCA-3’; *KALRN-CE* Reverse: 5’-GGACCAAGGTTCACAAATAGCAAAA-3’; *KLARN-CE* Reporter probe: 5’-AAGTGAGCAGCCATATTC-3’.

Relative expression of each cryptic exon was normalized to GAPDH using the DDCt method. For each differentiation batch, DCt values were first calculated by normalizing each sample’s Ct value to GAPDH, DDCt was then calculated by subtracting the average DCt of control samples within that batch from each sample’s DCt. Relative fold-change was calculated as 2^(-DDCt).

### RNA-seq data analysis and quality control

FASTQ files were aligned to hg38 human reference genome using STAR and reads were summarized with featureCounts. Raw gene-level counts were filtered to include only genes with expression above 1 count per million (CPM) in at least 90% of the samples. Raw read counts were converted to log_2_ CPM using the voom function from the limma R package.

#### Covariate assessment

To identify major sources of variation, principal component analysis (PCA) was performed on all genes using voom normalized expression values. For each co-variate (disease status: C9-ALS vs. control; differentiation batch; sex; hiPSC line; age; sequencing batch), a univariate linear model was fit against the first 10 PCs, and adjusted R2 values were calculated to determine variance contribution. Additionally, variancePartition was used to quantify covariate contribution to gene-level expression variability. A mixed model formula was applied (form <-∼ (1|disease) + (1|sex) + (1|patient) + (1|batch) + (1|seqrun) + (1|age)) and variance fractions were estimated using fitExtractVarPartModel.

#### Batch correction

Differentiation batch and sequencing batch were identified as key drivers of variation (Figures S1A, B). Because these two variables were highly correlated, both were included as covariates in removeBatchEffect from limma R package to generate a batch-corrected expression matrix for downstream WGCNA analysis.

#### Differential expression analysis

For each timepoint, differential expression analysis comparing C9-ALS and control samples was performed using a linear mixed-effects model. The design matrix was constructed with the following model: ∼ disease + sex + batch. The voom-normalized expression matrix was fit using lmFit. Empirical Bayes shrinkage of gene-wise variance estimate was applied using eBayes to stabilize variance. Differentially expressed genes (DEGs) were extracted using topTable and p-values adjusted for multiple testing using the Benjamini–Hochberg false discovery rate (FDR) method.

#### Alternative splicing analysis

Alternative splicing analysis was performed using Leafcutter following recommended parameters. Splice junctions were extracted from STAR-aligned RNA-seq bam files using regtools (Cotto et al., 2018) (1.0.0) ‘junctions extract’ using default parameters. Sample junctions were all clustered together using Leafcutter (Y. I. Li et al., 2018) script leafcutter_cluster_regtools.py to compare differentially spliced clusters between C9-ALS and control conditions. For differential splicing analysis between binary outcomes, leafcutter was run with script leafcutter_ds.R and exon files from GENCODE V31. Visualizations of splice graphs were constructed using leafviz pipeline. Percent spliced in (PSI) values were compared between C9-ALS and control samples for each timepoint. To assess differential splicing at the individual hiPSC line level, each C9-ALS line was compared against all control lines at each timepoint. To compare our hSpO differentially spliced events to those found in human datasets, we first filtered hSpO individual introns across all timepoints for significance (p < 0.01). For each human ALS dataset (Faura et al., 2025; O’Neill et al., 2025) the significant hSpO introns were matched to only retain introns that are found to be alternatively spliced in both human and hSpO. For each timepoint, a Pearson correlation was performed between the delta percent spliced in (ΔPSI) of hSpO and human introns, across all matched introns. Differentially spliced introns identified by Leafcutter (p < 0.05) in both the combined lines and individual lines per timepoint were compared to a reference list of known TDP-43 cryptic exon coordinates (Ma et al., 2022). For each known event, detection status and significance were recorded across all samples, and the percentage of known events recovered per sample was calculated.

### Weighted Gene Correlation Network Analysis (WGCNA)

WGCNA was performed using the blockwiseModules function with the following parameters: soft threshold power beta = 15.5, minModSize = 50, deepSplit = 3, and mergeCutHeight= 0.07. Hierarchical clustering of module members was performed to generate a dendrogram showing similar groupings of modules. The cutree function with k = 3 was applied to identify three main module clusters. Module correlation to disease state (C9 = 1, Control = 0) and age was determined with biweight midcorrelation (BiCor) analysis. Correlations were calculated for each timepoint (day 30, 50, 75) independently and across all samples irrespective of timepoint.

#### Module enrichment analysis

Fisher’s exact test (FET) was performed for each module’s members against three reference datasets:

1. (McEachin et al., 2025) proteomics dataset to determine module enrichment of proteins up or downregulated in C9orf72 postmortem spinal cords.
2. (Yadav et al., 2023) single cell RNA sequencing dataset of the human adult spinal cord to determine cell type-specific markers.
3. (Peters et al., 2025) single cell RNA sequencing dataset of day 60-80 hSpO to determine cell type-specific markers.

Genes and proteins with |log_2_fold-change|>1 and adj. p-value < 0.05 in reference datasets were considered for enrichment testing. Significance was determined as FET p-value <0.05.

*GO term enrichment*: Gene ontology (GO) Biological Processes enrichment was performed using clusterProfiler. Enriched terms passing Bonferroni-corrected p < 0.05 were retained. GO terms were manually grouped into predetermined functional categories based on biologically informative keywords in the GO term names for visualization.

#### Module eigengene analysis

To assess differences in module eigengene expression between control and C9-ALS, a t-test was performed for each timepoint independently in each module eigengene. Representative genes were chosen based on high correlation with module membership (kME values > 0.75).

### Nuclear-cytoplasmic fractionation

Nuclear and cytoplasmic fractions were isolated from hSpO using the Invitrogen PARIS™ kit (Invitrogen, AM1921). Five organoids per replicate were collected at each timepoint and all fractionations were performed on freshly collected organoids. Organoids were lysed in 150 µl of Cell Fractionation buffer to isolate cytoplasmic fractions, followed by lysis of the nuclear pellet in 150 µl of Cell Disruption buffer. Isolated fractions were then snap-frozen and stored at -80°C for downstream BCA analysis and Western blotting.

### Western blot

Protein concentration of lysed samples was determined using Pierce− BCA Protein Assay kit (Thermo Fisher, 23225) and samples were normalized to equal protein concentration. Samples were denatured using 2× Laemmli buffer (Biorad, 1610737) and boiled at 95°C for 5 minutes. Ten micrograms of samples were loaded onto 4-12% Bolt Bis-Tris Plus Gels (Thermo Fisher, NW04127BOX) and electrophoresed at 200V for 22 minutes using 1× Bolt− MES SDS running buffer (Thermo Fisher, B000202). Proteins were transferred onto nitrocellulose membranes (Thermo Fisher, IB23002) and blocked for 1 hour at room temperature in clear milk blocking buffer (Thermo Fisher, 37587). Primary (anti-TDP43: ProteinTech, 12892-1-AP, anti-acetyl-Histone H3: EMD Millipore, 06-599, anti-GAPDH: GeneTex, GTX627408) and secondary antibody solution (Dk anti-Mouse 488 AlexaFluor, Thermo Fisher Scientific A-21202; Dk anti-Rabbit 568 AlexaFluor, Thermo Fisher Scientific A10042) incubation was performed for 1 hour at room temperature with washes in 1× TBST (Thermo Fisher, 28360). Histone H3 served as a loading control for nuclear fractions, and GAPDH as a loading control for cytoplasmic fractions.

### DPR immunoassay

Two replicates per line, with two pooled organoids per replicate, were collected. Media was fully removed, washed with PBS, and organoids were snap frozen in dry ice. To each pellet, 50 µl of CoIP lysis buffer (50 mM TrisHCl, pH 7.4, 300 mM NaCl, 5mM EDTA, 1% Triton X-100) plus 10% SDS and 1× Halt Protease and Phosphatase Inhibitor Cocktail (Thermo Fisher Scientific, 78440) were added. Samples were lysed and sonicated. Prepared lysates were stored in -20°C until further processing. PolyGP dipeptide repeat (DPR) protein was quantified using a custom Meso Scale Discovery (MSD) immunoassay. All incubations (blocking, lysates, and antibody) were performed on an orbital shaker at 600 rpm, and all washes were performed using 150 µL of 0.05% TBS-Tween (TBS-T) between each incubation step. An affinity purified monoclonal poly-GP antibody (TALS 828.179; Target ALS) was both biotinylated and sulfo-taggedTM for capture and detection. Biotinylated poly(GP) antibody (30 µL,1 µg/mL) was immobilized onto a 96-well streptavidin plate and incubated overnight at 4°C without shaking. Plates were then blocked using MSD blocking buffer A for 1 hour at room temperature. hSpO lysates were diluted to 0.6 µg/mL, and 50 µL of lysate was loaded in technical duplicates for 2 hours at room temperature. Sulfo-tagged™ poly(GP) antibody (30 µL, 1 µg/mL) was then added into the same wells and incubated for 1.5 hours at room temperature. Following incubation, 150 µL of 2× Read buffer T (R92TC-3, MSD) was added to each well and plates were read on MSD QUICKPLEX SQ210.

### Quantitative proteomics

Three organoids per replicate were placed into RINO microcentrifuge tubes (Next Advance, TUBE1R5S), media removed, and samples snap frozen on dry ice for storage at -80°C until further processing.

#### Protein Extraction, Digestion, and DIA-MS Sample Preparation

hSpO samples were homogenized in 8 M urea lysis buffer (Sigma, U0631) supplemented with 1× Halt Protease Inhibitor Cocktail and 1× Halt Phosphatase Inhibitor Cocktail (Thermo Fisher Scientific, 78440). Samples were disrupted using stainless-steel beads (Next Advance, Inc., SSB14B) and homogenized with a BeadBlaster homogenizer (Benchmark Scientific) for two cycles of 5 minutes at speed 12. Following homogenization, lysates were subjected to probe sonication (25% amplitude; 5 s pulse, 5 s rest, 3 cycles) and centrifuged at 13,300 rpm for 10 minutes to remove insoluble material. Protein concentration was determined using the Pierce− BCA Protein Assay Kit (Thermo Fisher Scientific, 23225). Lysates were normalized to 2 mg/mL prior to proteomic analysis.

For digestion, 100 µg of total protein was reduced with 50 mM dithiothreitol (DTT; Sigma, D5545) for 30 minutes at room temperature with agitation (800 rpm) and subsequently alkylated with 100 mM iodoacetamide (IAA; Sigma, I1149) for 30 minutes in the dark at room temperature. Samples were diluted with 100 mM Tris-HCl (pH 8.0) to reduce the urea concentration below 4 M and digested overnight at room temperature using endoproteinase LysC (Wako Chemicals, 125-05061) at an enzyme-to-substrate ratio of 1:15 (w/w). Samples were further diluted with 100 mM Tris-HCl (pH 8.0) to reduce the urea concentration below 1 M and digested overnight with sequencing-grade modified trypsin (Promega) at an enzyme-to-substrate ratio of 1:15 (w/w).

Digestion was terminated by acidification with an equal volume of 4% phosphoric acid (Sigma, PX0996-6) and incubated at room temperature for approximately 20 minutes with agitation. Peptides were subsequently purified using Oasis® MCX Prime 10 mg µElution plates (Waters Corporation) according to a mixed-mode cation-exchange workflow. Samples were loaded onto MCX cartridges and washed sequentially with 100 mM ammonium formate containing 2% formic acid followed by methanol. Peptides were eluted using 5% ammonium hydroxide (ThermoFisher Scientific, 460801000) in methanol, and dried to completeness in a SpeedVac vacuum concentrator (Thermo Fisher Scientific). Approximately 25 µg of purified peptides from each sample were used for data-independent acquisition mass spectrometry (DIA-MS).

#### DIA-MS Proteomics

Proteomic profiling was performed using an unbiased data-independent acquisition mass spectrometry (DIA-MS) approach. All samples (500 ng in 20 µL per sample) were were loaded and separated using a Bruker timsTOF HT mass spectrometer coupled to an Evosep One liquid chromatography system. Samples were acquired at 30 samples per day (SPD) using a Performance Column (8 cm × 150 µm × 1.5 µm; part number EV1109). DIA acquisition was performed using a narrow ion mobility window (0.85–1.35 V·s/cm^2^). Database searches and protein quantification were performed on raw files using DIA-NN 1.9.1 (Data-Independent Acquisition by Neural Networks) (Demichev et al., 2020) with default, fully tryptic search parameters (raw data provided in Table S1L). MS/MS spectra were searched against all canonical human proteins downloaded from UniProt (20,383 entries), along with decoy sequences (47,262 entries). Precursor mass tolerance was set to the recommended MS1 mass accuracy setting per raw data file. Enzyme specificity was set to strict trypsin, allowing up to one missed cleavages Peptide length was restricted to 7–30 residues, and peptide mass to 300–1,800 m/z. The FDR threshold was set to 1%.

#### Bioinformatics processing and statistical analysis

A Tunable Approach for Median Polish of Ratios (TAMPOR), mode 4, was applied to DIA-MS data (Dammer et al., 2023), and peptides or proteins missing in ≥50% of samples were removed, as previously described (Johnson et al., 2020, 2022). Multidimensional scaling was used to visualize the distribution of post-TAMPOR data as a quality control step. Fast parallelized one-way ANOVA with Benjamini-Hochberg correction for multiple comparisons was performed within each disease group using an in-house script (https://github.com/edammer/parANOVA) to identify differentially expressed proteins (Table S1N). Differential abundance was visualized using volcano plots generated with the ggplot2 package (Wickham, 2016).

### Whole organoid immunohistochemistry

Whole organoids were fixed in 4% paraformaldehyde (PFA) in PBS for 2 hours at 4°C. Organoids were placed in blocking solution (10% normal donkey serum - Sigma S30-M, 0.3% Triton X-100 diluted in PBS) for 4 hours, followed by primary antibody solution (blocking solution without Triton) for 2–3 days (anti-ISL1: R&D Systems, AF1837). Organoids were then washed three times for 30 minutes per wash, then placed in secondary antibody solution (blocking solution without Triton) overnight protected from light (Dk anti-Goat 647 AlexaFluor: Invitrogen, A-21447). Organoids were washed once for 30 minutes, placed in Hoechst solution (1:10,000 Fisher Scientific, H3569) for 20-30 minutes, then washed a final time for 30 minutes before being placed on slides with coverslips (Fisher Scientific, S24461) using Aquamount (Polysciences, 18606-20) in 3-5 stacked iSpacers (SunJin Lab, IS213) per slide.

### Organoid cryopreservation and immunohistochemistry

Organoids were fixed in 4% PFA (Electron Microscopy Sciences 15710) for 2 hours. Fixation was followed by three PBS washes. Organoids were placed in 30% sucrose solution in PBS for 24–48 hours or until samples sank to the bottom, embedded in 1:1 30% sucrose:OCT solution (Tissue-Tek OCT Compound, 4583) in molds (Fisher, 22-363553), and frozen at -80°C. For immunohistochemistry, 15 µm-thick sections were cut using a cryostat (Leica) onto slides (VWR, 48311-703). PAP pen (Fisher Scientific, NC0822416) was used to create hydrophobic regions around cryosections. Cryosections were then washed with PBS to remove excess OCT, blocked for 1 hour at room temperature (10% normal donkey serum, Sigma, S30-M, 0.3% Triton X-100 diluted in PBS), and incubated overnight at 4°C with primary antibodies in blocking solution without Triton (anti-CHAT: Millipore, AB144P, anti-VGLUT2: Synaptic Systems, 135403, anti-GABA: Sigma, A2052, anti-NFH: Abcam ab4680, anti-MAP2: SySy 188004). The next day, cryosections were washed with PBS and then incubated with secondary antibodies in blocking solution without Triton for 1 hour at room temperature protected from light (Dk anti-Chicken 488 AlexaFluor, Jackson, 703-545-155; Dk anti-Guinea Pig 647 AlexaFluor, Jackson, 706-605-148; Dk anti-Goat 488 AlexaFluor, Abcam, ab150133; Dk anti-Chicken 790 AlexaFluor (AffiniPure), Jackson, 703-655-155, Dk anti-Rabbit 568 AlexaFluor, Invitrogen, A10042; Dk anti-Rabbit 647 AlexaFluor, Invitrogen, A78957). Following washes with PBS, nuclei were visualized with Hoechst 33258 (1:10,000 applied for 5 minutes, Fisher Scientific, H3569). Finally, slides were mounted for microscopy with cover glasses (Fisher Scientific, S24461) using Aquamount solution (Polysciences, 18606-20). For TDP-43 specific immunohistochemistry, organoids were sectioned at 10 µm, blocked for 1 hour at room temperature (5% bovine serum albumin in 0.1% Triton X-100 diluted in PBS), incubated in primary antibody solution for 1 hour at room temperature (anti-TDP43: ProteinTech, 12892-1-AP), and incubated in secondary antibody solution for 1 hour at room temperature (Dk anti-Rabbit 568 AlexaFluor, Invitrogen, A10042). All other washing and mounting steps were performed as above.

### Fluorescence in-situ hybridization (FISH)

Organoid samples were prepared as described above. Thawed sections were washed once with Dulbecco’s phosphate-buffered saline (DPBS, Gibco, 14190250) before fixation with 4% Paraformaldehyde for 15 minutes, then washed again, and permeabilized with 0.2% Triton X-100 in 1× PBS for 10 minutes. After a third wash, samples were incubated with 2× Saline sodium citrate (SSC, VWR, 10128-690) for 10 minutes. Hybridization buffer was prepared on ice by combining 20× SSC, Sodium Phosphate (1M, pH 7.0, VWR, BDH9298-500G), 33.3% Dextran Sulphate (Millipore Sigma, D8906-10G), and 20× Vanadyl Complex (Sigma-Aldrich, R3380-5ML) with formamide (VWR, IB72024) added immediately before use. The probe (/5TYE563/CCCCGGCCCCGGCCCC, Qiagen) was diluted in 100 µl of hybridization buffer, denatured at 95°C for 5 minutes and immediately transferred to ice. The denatured probe was then combined with the remaining hybridization buffer and added to slides to incubate for 2 hours at 66oC in a humidified chamber. Following hybridization, slides were washed twice in 2× SSC with 0.1% Tween-20 for 10 minutes at room temperature, followed by a single wash in 0.1× SSC for 10 minutes, and a final rinse in deionized water. Slides were washed in Hoechst solution (1:10,000 applied for 5 minutes, Fisher Scientific, H3569) then washed a final time in deionized water. Coverslips were mounted using Aquamount (Polysciences, 18606-20).

### FISH imaging and quantification

For each organoid, one field of view image was taken in a total of three sections spanning the width of the whole organoid. Images were taken on a Leica Stellaris 5 confocal microscope using a 63× objective. The percentage of nuclei containing RNA foci was quantified using a custom analysis pipeline in ImageJ. Raw images were first converted to maximum intensity projections. The nuclear channel (Hoechst) was segmented as follows: Make Binary > Watershed > Fill Holes. Nuclear regions of interest were identified using Analyze Particles with a size threshold of 100–infinity pixels and circularity of 0–1. RNA foci were detected using Find Maxima with a prominence threshold that was adjusted for each paired control/C9-ALS slide to restrict output to single points and minimize background. Several control/C9-ALS slides were visually inspected after thresholding to ensure that automatic quantification was detecting all visible foci and minimal debris. All fluorescence measurements were exported as CSV for downstream analyses. Foci counts per nucleus were calculated by dividing the RawIntDen value for each nuclear ROI by 255 to yield the discrete number of foci. Samples were classified as foci+ if they contained more than one focus. For each sample, total nuclear count, foci+ nuclei count, and percentage of foci+ nuclei were calculated.

### TDP-43 nuclear intensity quantification

Following immunohistochemistry as described above, cryosections were imaged on Keyence BZ-X810 fluorescence microscope using a 60× immersion objective with one region of interest (ROI) per section and three sections per hSpO. Regions were chosen unbiasedly by picking areas with healthy nuclei present and TDP-43 channel turned off. Multi-channel z-stacks were processed in ImageJ/Fiji. Nuclear ROIs were defined with a custom script using StarDist2D. Briefly, the Hoechst channel was processed with StarDist2D using the “Versatile (fluorescent nuclei)” pre-trained model. Established nuclear ROIs were used to extract mean gray-scale intensity values from the TDP-43 channel. Across all cells detected in all sections per organoid, the median nuclear TDP-43 intensity was calculated to generate one value per organoid.

### ISL1 cell counting

hSpO were fixed with 4% PFA and immunohistochemistry performed as described above. Imaging was performed on a Leica Stellaris 5 confocal microscope with z-step size of 15 µm across 12-15 z-steps to capture the entire hSpO volume. Quantification of ISL1+ cells was performed using a custom ImageJ script. The macro workflow was comprised of the following steps: (1) First, a Gaussian blur was applied (sigma = 3) to smooth noise, (2) then the blurred image was subtracted from the original to enhance contrast, (3) followed by particle size threshold set between 48 and 255 to isolate ISL1+ cell bodies, (4) application of a binary mask, (5) and watershed segmentation to allow individual cell counting. Finally (6) particle analysis was performed to count objects within size range 20-500 pixels. The total number of ISL1+ cells per organoid was recorded for each timepoint.

### Lactate Dehydrogenase (LDH) Assay

Cytotoxicity was assessed by measuring lactate dehydrogenase (LDH) release into conditioned media using the LDH-Glo− Cytotoxicity Assay (Promega; J2381), according to the manufacturer’s instructions. Organoids were cultured in 24-well plates (VWR; 62406-183), with two organoids per well and two wells per hiPSC line. Conditioned media was collected weekly prior to replacement with 1 mL of fresh hSpO media. Collected media samples were diluted 1:9 in LDH storage buffer (200 mM Tris-HCl, 10% glycerol, 1% bovine serum albumin), snap frozen on dry ice and stored at −80°C. Maximum LDH release (positive control) was determined by treating 1-2 organoids with 0.2% Triton X-100 for 20 minutes at 37°C. Background LDH levels were determined using fresh hSpO media, which was snap frozen and stored at −80°C prior to assay measurement. Equal volumes of LDH Detection Reagent were added to each sample in an opaque 96-well plate (Corning, 3912). Luminescence was measured after 60 minutes of incubation at room temperature in the dark using a microplate reader (FLUOstar Omega; BMG Labtech). Percent cytotoxicity was calculated using the following formula:

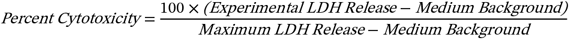

### CRIP analysis

Differential splicing was detected using LeafCutter as described above. Splicing-altered introns for CRIP analysis (Group A) were defined as introns with |ΔPSI| > 0.2 and adj. P-value < 0.05. Genomic coordinates were lifted from hg38 to hg19 using UCSC liftOver, retaining 35, 39, and 53 introns at days 30, 50, and 75, respectively. For each timepoint, a background set (Group B) was generated by randomly sampling introns with minimal evidence of differential usage (adj. p-val > 0.9), using a 5:1 ratio to Group A. For each intron in Group A or Group B, the 400-nucleotide (nt) genomic sequences flanking the 5′ (upstream) and 3′ (downstream) splice sites were extracted in a strand-aware manner, such that upstream and downstream were defined relative to the transcriptional orientation of each intron. The 5′ and 3′ splice-site flanking regions were analyzed independently throughout the study.

Sequence-based RBP-binding prediction was performed with CRIP (Zhang et al., 2019), a deep-learning model that predicts RBP-binding probability from RNA sequence. Each 400-nt flank was tiled into 300 overlapping 101-nt windows, and each window was scored against 137 RBPs to generate per-window RBP-binding probabilities (0-1) for each RBP. For each combination of intron, flank, and RBP, the 300 per-window probabilities were summarized into two metrics: peak count, defined as the number of windows with binding probability > 0.8; and binding intensity, defined as the sum of the window probabilities exceeding 0.5. Both metrics were computed separately for the statistics of upstream and downstream flanks. Analyses were performed separately for each timepoint (days 30, 50, 75), binding metric (peak count, binding intensity), and flank (upstream, downstream). Within each stratum, each of the 137 RBPs was tested for higher predicted binding in Group A than in Group B using a one-sided Wilcoxon rank-sum (Mann-Whitney U) test. Effect sizes were quantified as Cliff’s delta (Fiel Peres, 2026). Positive values (range -1 to +1) indicate higher predicted RBP binding in Group A compared with Group B. Within each stratum, p-values across the 137 RBPs were corrected for multiple testing using the Benjamini-Hochberg procedure. RBPs were considered nominally significant at p ≤ 0.05 with higher predicted binding in Group A.

Both nominal and FDR-adjusted p-values, group summary statistics, and effect sizes are reported in **Tables S1T–V**.

## Data and code availability

Feature counts from bulk RNA sequencing data generated in this study will be made available upon publication. All code is available from the lead contact upon request.

## Protocol availability

The Emory University Brain Organoid Hub generated and maintained many of the hiPSC lines and organoid differentiations used for these experiments. Detailed protocols and demonstration videos on hiPSC thawing, maintenance, passaging, and organoid differentiation are available at www.brainorganoidhub.com/protocols.

## Statistical analysis

GraphPad Prism was used for statistical analysis of immunohistochemistry, RNA foci, immunoassay, LDH assay, Western blotting, and RT-qPCR experiments. All sequencing data, differential splicing analysis, and ISL1 counting statistics were performed in R. All data underwent normality testing prior to selecting appropriate statistical tests. Figure legends detail exact statistical tests used for each experiment. Data represent mean ± SD unless otherwise noted. Statistical significance is denoted as p < 0.05 (*), p < 0.01 (**), and p < 0.001 (***) unless otherwise noted.

---

**Figure S1:**
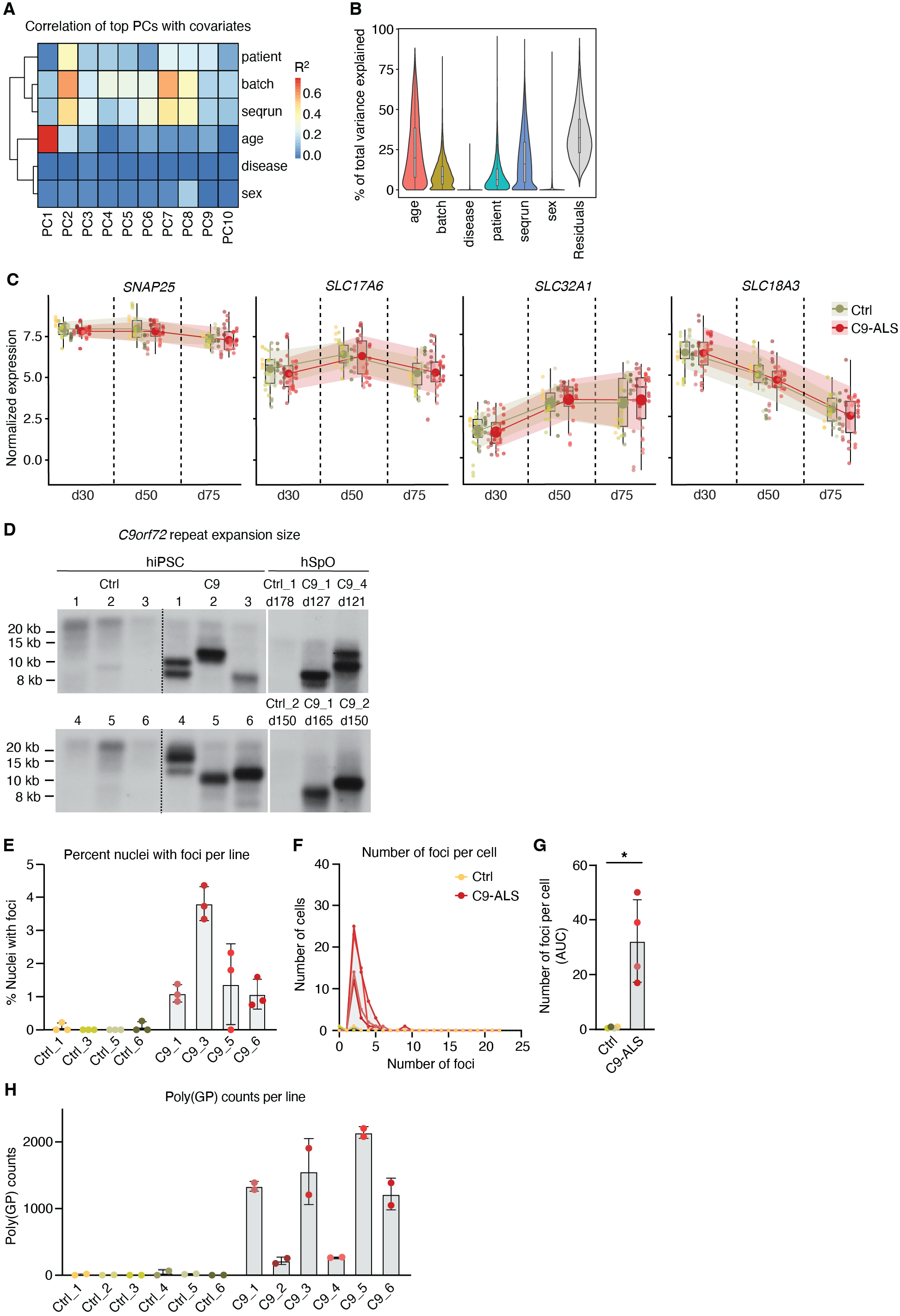
C9orf72 relevant pathologies prevalence in C9-ALS hSpO, related to Figure 1. **(A)** Heatmap showing correlation of each covariate to principal components (PCs), colored by R2 value. **(B)** Violin plot showing the percentage of each covariate that contributes to variation in gene expression in bulk RNA sequencing dataset. **(C)** Box plots showing trimmed mean of M-values (TMM) normalized expression for neuronal cell-type markers over time in C9-ALS and control hSpO. Each dot represents a sample from **Figure 1C**, colored by the hiPSC line they were derived from. Middle black lines represent median and colored large dots represent mean expression in control (green) and C9-ALS (red) hSpO. Shaded green and red regions represent standard deviation in control and C9-ALS conditions, respectively. **(D)** Southern blot analysis of the G_4_C_2_ repeat in control and C9-ALS hiPSCs and organoids. **(E)** Bar plot showing percent of nuclei with G4C2+ RNA foci in control and C9-ALS hSpO at day 75, separated per hiPSC line. Each dot represents an average number of foci per organoid, with 3 cryosections and one field of view per section counted per organoid. n = 3 organoids per line, with n = 4 hiPSC lines in each control and C9-ALS condition. **(F)** Histogram showing the distribution of the number of foci per cell colored by hiPSC line. n = 3 organoids per line, with n = 4 hiPSC lines in each control and C9-ALS condition. n cells with foci (range indicating numbers per section quantified) = 3–10 (C9_1), 1–14 (C9_3), 5–19 (C9_5), 1–8 (C9_6). n foci per cell = 2–6 (C9_1), 2–9 (C9_3), 2–5 (C9_5), 2–5 (C9_6). **(G)** Bar plot showing area under the curve (AUC) quantification for histogram in F. Each dot represents an hiPSC line, with 3 organoids per line and 3 sections per organoid. Mann-Whitney test (*p = 0.028). **(H)** Bar plot showing poly(GP) counts in day 75 control and C9-ALS hSpO separated by line. n = 2 biological replicates per hiPSC, with n = 4 hiPSC lines in each control and C9-ALS condition. Each replicate represents 2 organoids pooled together.

**Figure S2:**
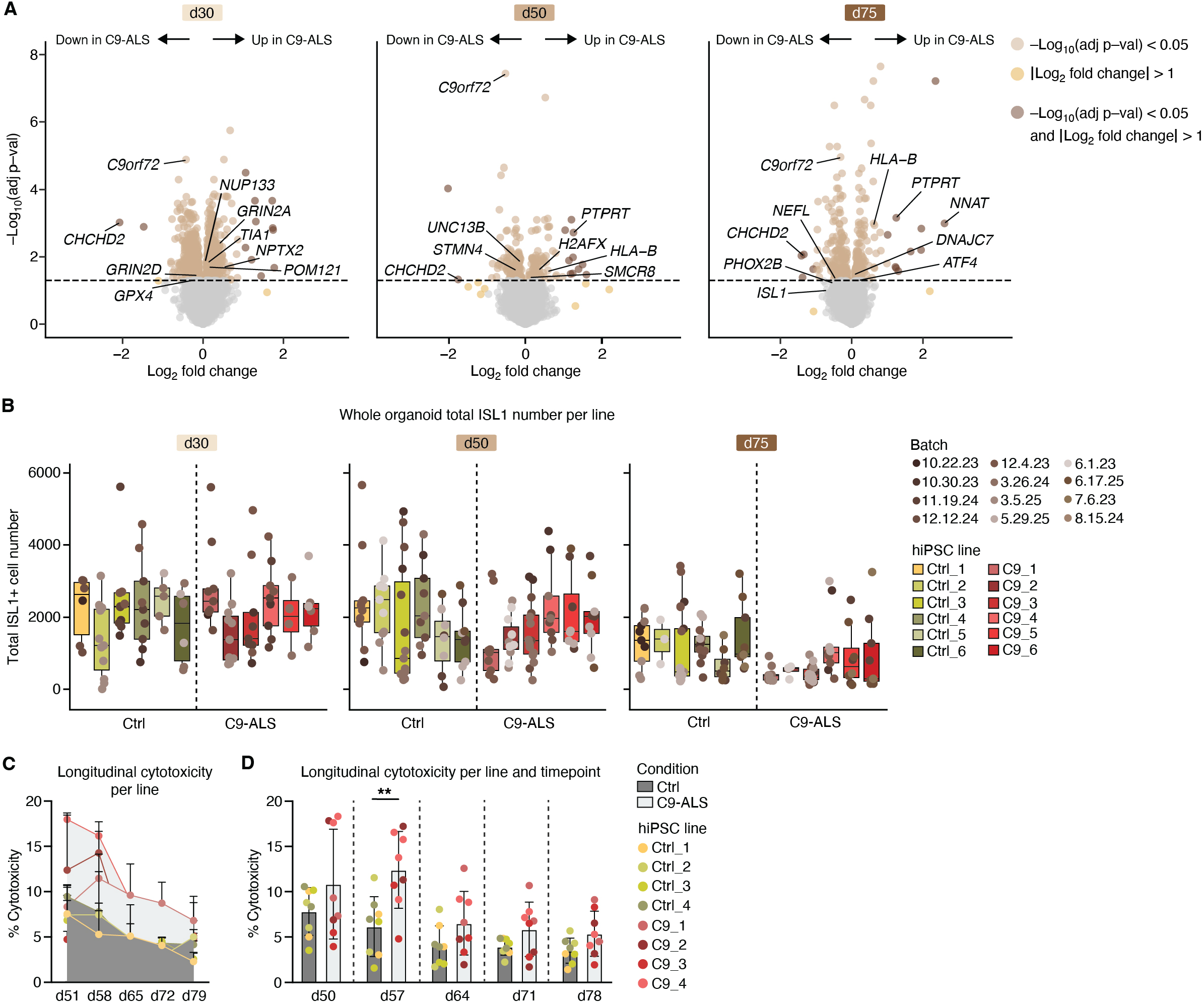
Neuronal dysfunction and cell death in C9-ALS hSpO, related to Figure 1. **(A)** Volcano plots showing differential gene expression in C9-ALS vs control hSpO at days 30, 50 and 75. **(B)** Bar plot showing distribution of total ISL1^+^ counts across days 30, 50 and 75 separated by line and colored by differentiation batch. Each dot represents an individual organoid. n = 6 hiPSC lines in each control and C9-ALS condition. **(C)** Line graph showing percent cytotoxicity in control and C9-ALS hSpO over time measured by lactate dehydrogenase (LDH) assay. Each dot per timepoint represents an average of 2 replicates. Each replicate was collected from a well of a 24-well plate with x2 hSpO each. Shaded regions indicate control or C9-ALS hiPSC lines. **(D)** Bar plot showing percent cytotoxicity in control and C9-ALS hSpO over time. Each dot represents a replicate. Each replicate was collected from a well of a 24-well plate with x2 hSpO each. n = 2 replicates per hiPSC line, with n = 4 hiPSC lines in each control and C9-ALS condition. Mixed effects analysis with Day (F(1.98, 27.75) = 12.76, ***p = 0.0001) and Condition (control vs C9-ALS; F(1, 14) = 8.163, *p = 0.013) as factors (Condition × Day interaction, F(4, 56) = 1.76, p = 0.15), followed by Tukey’s multiple comparison test per timepoint (day 50: p = 0.23, day 57: **p = 0.006, day 64: p = 0.12, day 71: p = 0.12, day 78: p = 0.09).

**Figure S3:**
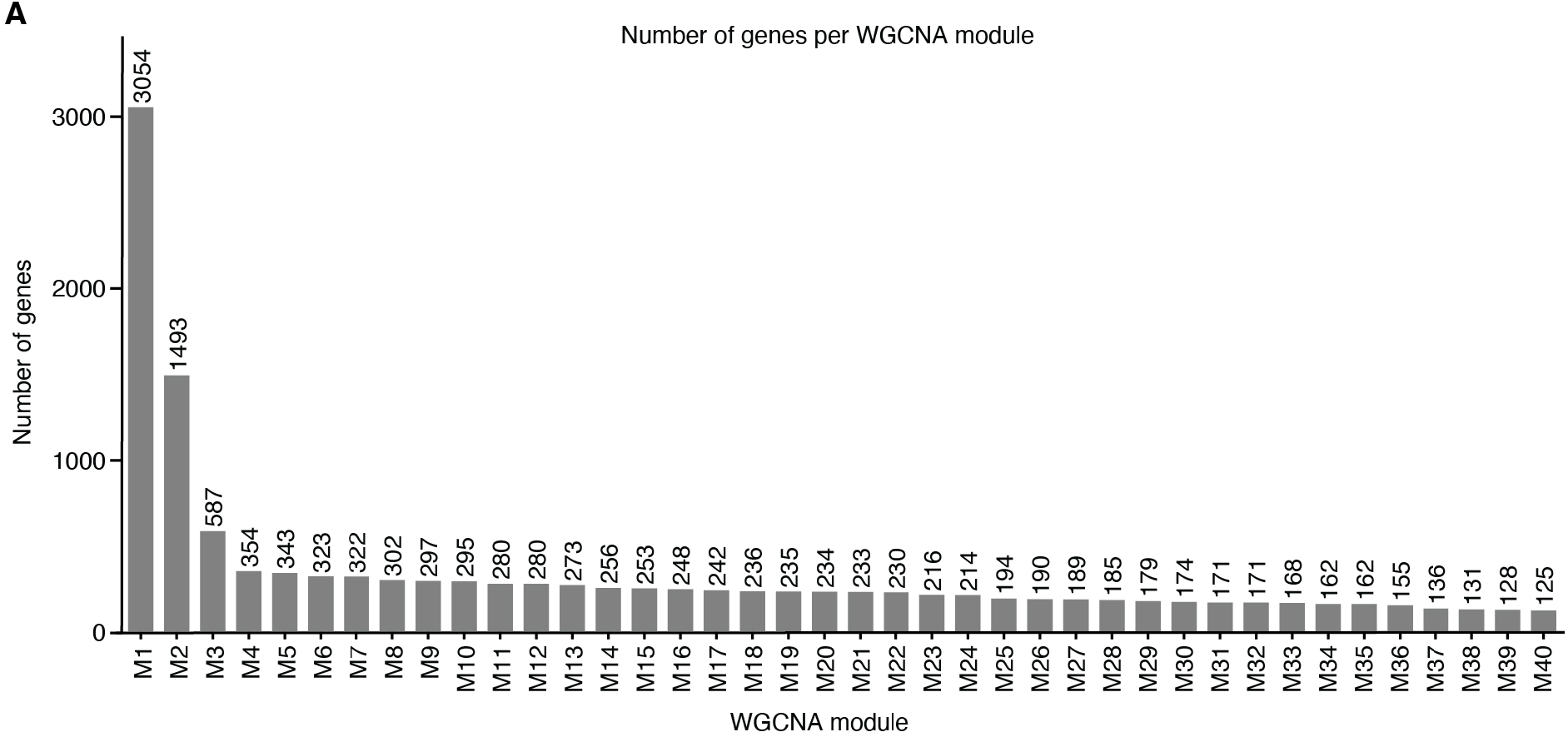
Number of genes per module, related to Figure 2. **(A)** Bar plot showing the number of genes in each WGCNA module.

**Figure S4:**
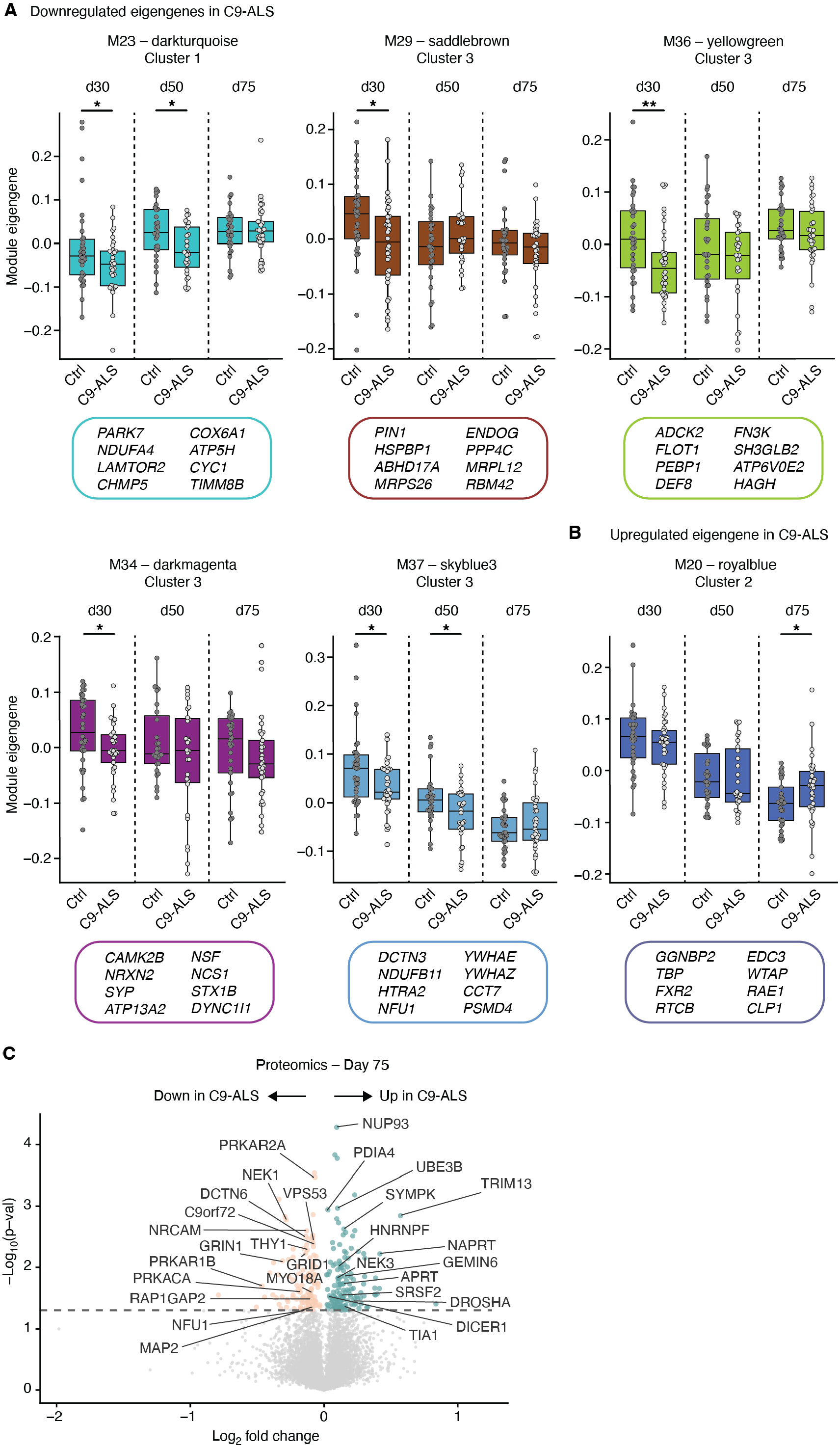
Module eigengenes and proteomics show dysregulated processes over time in C9-ALS hSpO, related to Figure 3. **(A)** Boxplots showing module eigengenes that are decreased in C9-ALS hSpO from clusters 1 and 3. Boxes represent the median and interquartile range (IQR: 25^th^-75^th^ percentile), with whiskers extending to 1.5 × IQR. Example genes per module are shown in corresponding colored boxes below. Each dot represents a sample from **Figure 1C**. T-tests were performed for each timepoint independently (*p < 0.05, **p < 0.01). **(B)** Boxplots showing module eigengene that is increased in C9-ALS hSpO for a representative module from cluster 2. Boxes represent the median and interquartile range (IQR: 25^th^-75^th^ percentile), with whiskers extending to 1.5 × IQR. Example genes per module are shown in corresponding colored boxes below. Each dot represents a sample from **Figure 1C**. T-tests were performed for each timepoint independently (*p < 0.05). **(C)** Volcano plot showing upregulated (teal) and downregulated (salmon) proteins in C9-ALS from day 75 hSpO proteomics. Highlighted dots show proteins with p < 0.05 and |Log_2_ Fold Change| > 0.

**Figure S5:**
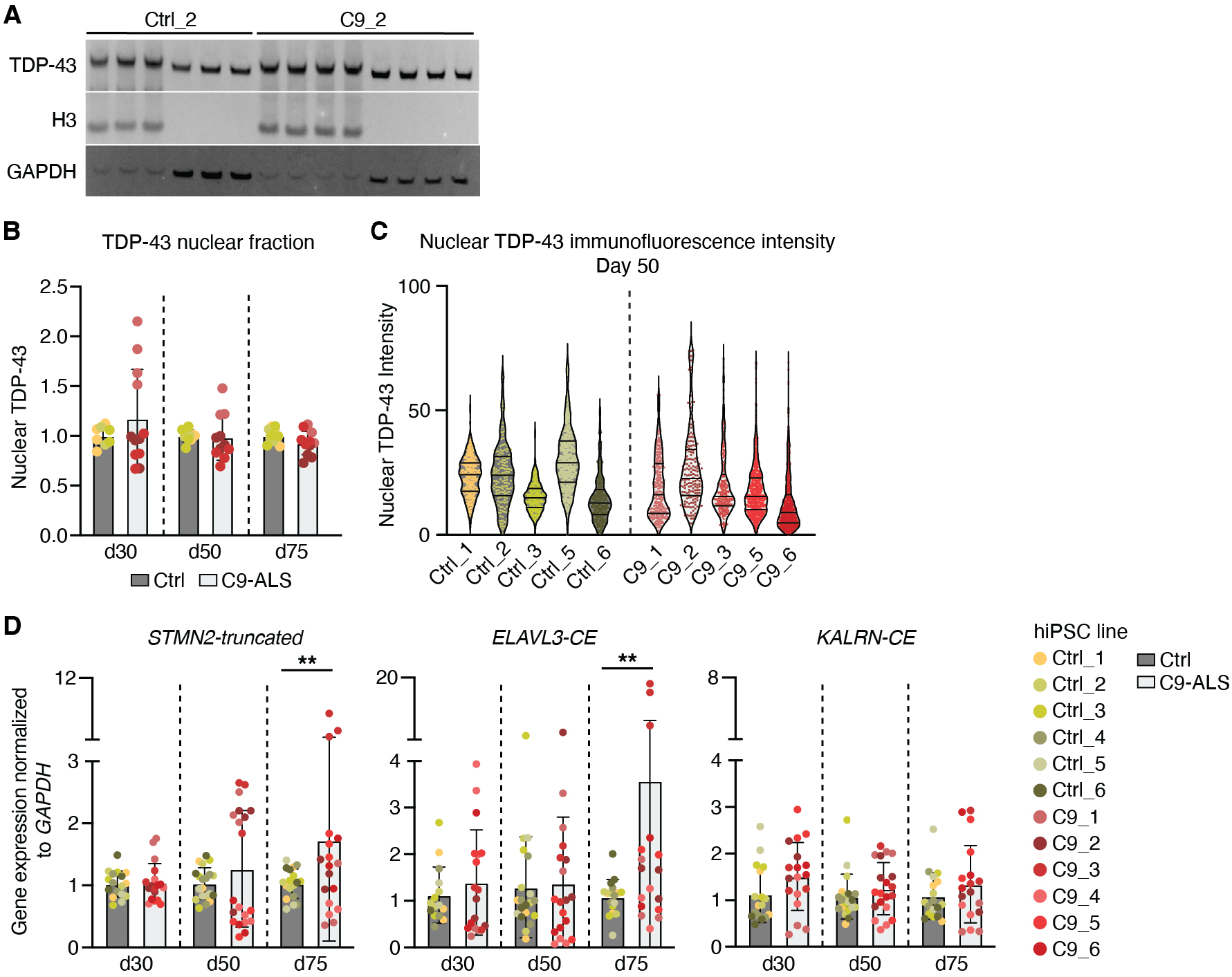
Nuclear TDP-43 levels and prevalence of cryptic exons in C9-ALS hSpO, related to Figure 4. **(A)** Representative image of Western blot for TDP-43, H3 (nuclear control), and GAPDH (cytoplasmic control), demonstrating clean isolation of subcellular fractions. **(B)** Bar plot showing quantification of nuclear TDP-43 Western blots in hSpO nuclear fractions over time. Each dot represents a replicate, with 5 organoids pooled and fractionated per replicate. n = 2-4 replicates per hiPSC line, with n = 3 hiPSC lines in each control and C9-ALS condition. All C9-ALS samples were normalized to batch controls per timepoint. Mixed effects analysis with Day (F(1.313, 24.30) = 1.940, p = 0.17) and Condition (control vs C9-ALS; F(1, 21) = 0.2088, p = 0.65) as factors (Day × Condition interaction, F(1.313, 24.30) = 2.132, p = 0.15), followed by Tukey’s multiple comparisons test per timepoint (day 30: adj. p-val = 0.26, day 50: adj. p-val = 0.85, day 75: adj. p-val = 0.09). **(C)** Violin plots showing nuclear TDP-43 fluorescence intensity in individual hSpO cells at day 50. Each dot represents the intensity of one cell in a hSpO cryosection, colored by the hiPSC line they were derived from, with a total of 3 organoids per hiPSC lines, n = 5 hiPSC lines in each control and C9-ALS condition. For each organoid, three cryosections were imaged and quantified with the following number of cells counted per line: n = 776 (Ctrl_1), 318 (Ctrl_2), 666 (Ctrl_3), 1,499 (Ctrl_5), 631 (Ctrl_6), 680 (C9_1), 396 (C9_2), 469 (C9_3), 1,232 (C9_5), 1,108 (C9_6). **(D)** RT-qPCR barplots showing expression of *STMN2, ELAVL3* and *KALRN* cryptic exons normalized to *GAPDH*. Each dot represents a replicate per hiPSC line, with 2-3 organoids pooled per replicate, n = 3-4 biological replicates per line, and n = 6 hiPSC lines in each control and C9-ALS condition. All C9-ALS samples were normalized to batch controls at each timepoint. Mixed effects analysis with Day and Condition (control vs C9-ALS) as factors for each cryptic exon independently, followed by Tukey’s multiple comparisons test at each timepoint. *STMN2*-*truncated*: Day (F(2, 67) = 2.23, p = 0.16); Condition (F(1, 38) = 3.17, p = 0.08); Day × Condition interaction (F(2, 67) = 2.18, p = 0.12); day 30, adj. p-val = 0.95; day 50, adj. p-val = 0.34; day 75, ***adj. p-val = 0.009. *ELAVL3-CE*: Day (F(2, 100) = 2.18, p = 0.12); Condition (F(1, 100) = 4.23, *adj. p-val = 0.04); Day × Condition interaction (F(2, 100) = 2.80, p = 0.06); day 30, adj. p-val = 0.74; day 50, adj. p-val = 0.92; day 75, **adj. p-val = 0.003. *KALRN-CE*: Day (F(2, 67) = 0.82, p = 0.44); Condition (F(1, 38) = 3.46, p = 0.07); Day × Condition interaction (F(2, 67) = 0.28, p = 0.76); day 30, adj. p-val = 0.09; day 50, adj. p-val = 0.39; day 75, adj. p-val = 0.23.

**Figure S6:**
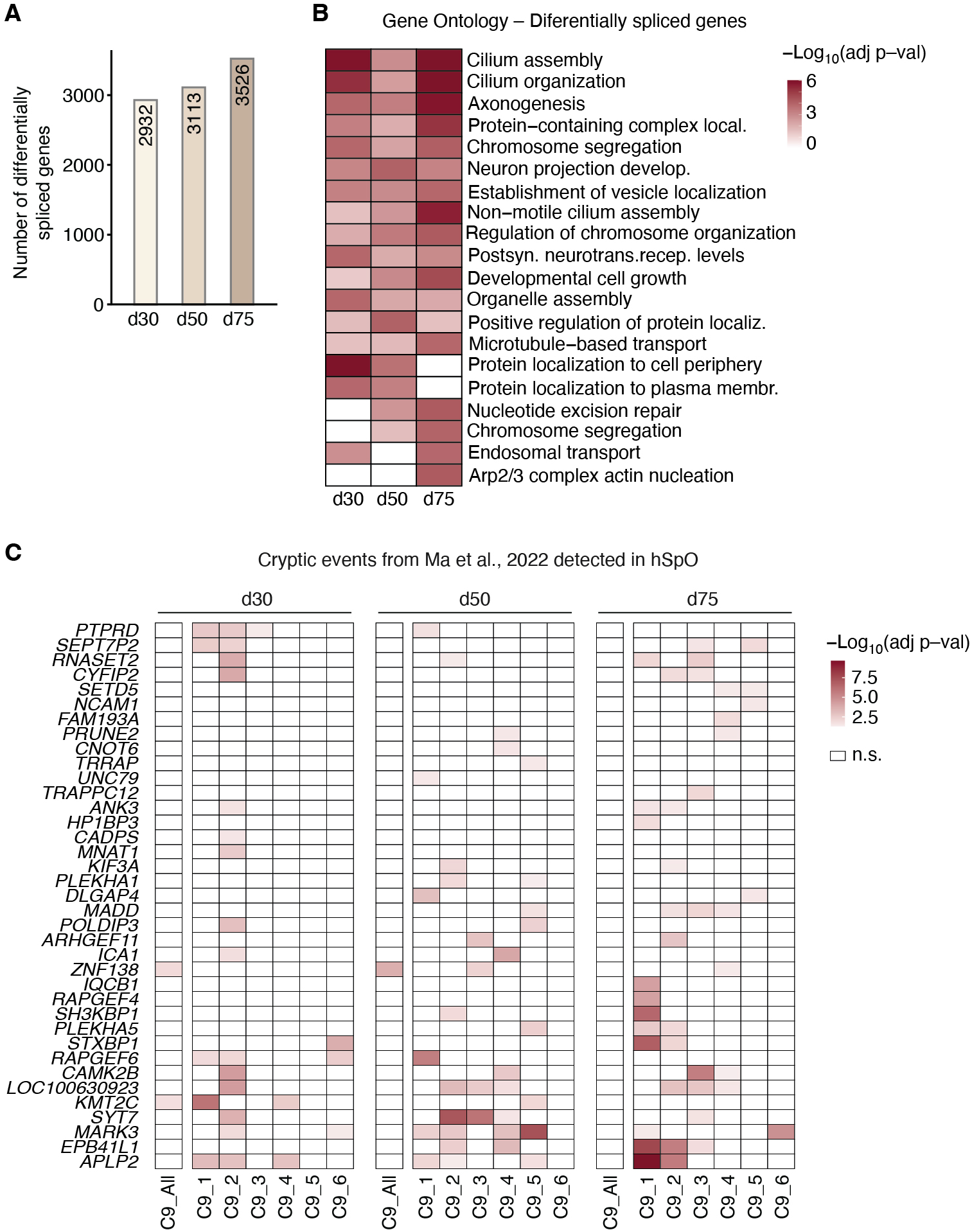
Differential splicing analysis and TDP-43 relevant splice targets in C9-ALS, related to Figure 4. **(A)** Bar plot showing the number of significant differentially spliced genes in hSpO over time from Leafcutter analysis. **(B)** Heatmap showing gene ontology (biological processes) terms enriched among differentially spliced genes at each timepoint colored by -log_10_(p-value). **(C)** Heatmap showing known cryptic events from Ma et al., 2022 in hSpO per timepoint from either combined analysis (C9_all) or per-line Leafcutter analysis. Color indicates -log_10_(p-value). Only splicing events from Ma et al., 2022 that were significantly differentially spliced in at least one hiPSC line and/or timepoint are shown.

**Figure S7:**
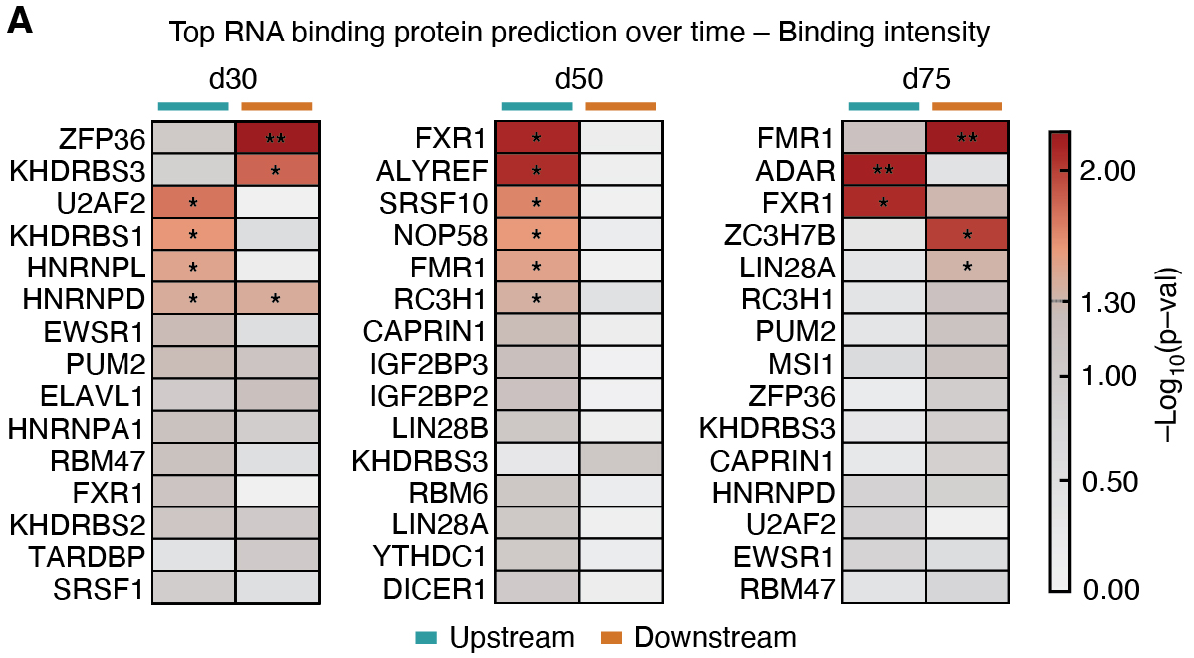
RNA binding protein predictions from most significant splice events, related to Figure 5. **(A)** Heatmap showing top predicted RBPs per timepoint based on binding intensity metric and colored by p-value from Wilcoxon rank sum test comparing binding intensity per RBP in altered vs unaltered group of splicing events defined in **Figure 5A** (*p<0.05). Dotted line in scale bar legend indicates significance threshold (−log_10_(0.05) = 1.30). Values for both upstream and downstream flanks of splicing event are separately shown.

